# Erg251 has complex and pleiotropic effects on azole susceptibility, filamentation, and stress response phenotypes

**DOI:** 10.1101/2024.03.06.583770

**Authors:** Xin Zhou, Audrey Hilk, Norma V. Solis, Bode M. Hogan, Tessa A. Bierbaum, Scott G. Filler, Laura S. Burrack, Anna Selmecki

**Affiliations:** Department of Microbiology and Immunology, University of Minnesota, Minneapolis, MN, USA; Division of Infectious Diseases, Lundquist Institute for Biomedical Innovation at Harbor UCLA Medical Center, Torrance, CA, USA; Gustavus Adolphus College, Department of Biology, Saint Peter, MN, USA; David Geffen School of Medicine at UCLA, Los Angeles, CA, USA

## Abstract

Ergosterol is essential for fungal cell membrane integrity and growth, and numerous antifungal drugs target ergosterol. Inactivation or modification of ergosterol biosynthetic genes can lead to changes in antifungal drug susceptibility, filamentation and stress response. Here, we found that the ergosterol biosynthesis gene *ERG251* is a hotspot for point mutations during adaptation to antifungal drug stress within two distinct genetic backgrounds of *Candida albicans*. Heterozygous point mutations led to single allele dysfunction of *ERG251* and resulted in azole tolerance in both genetic backgrounds. This is the first known example of point mutations causing azole tolerance in *C. albicans.* Importantly, single allele dysfunction of *ERG251* in combination with recurrent chromosome aneuploidies resulted in *bona fide* azole resistance. Homozygous deletions of *ERG251* caused increased fitness in low concentrations of fluconazole and decreased fitness in rich medium, especially at low initial cell density. Dysfunction of *ERG251* resulted in transcriptional upregulation of the alternate sterol biosynthesis pathway and *ZRT2*, a Zinc transporter. Notably, we determined that overexpression of *ZRT2* is sufficient to increase azole tolerance in *C. albicans*. Our combined transcriptional and phenotypic analyses revealed the pleiotropic effects of *ERG251* on stress responses including cell wall, osmotic and oxidative stress. Interestingly, while loss of either allele of *ERG251* resulted in similar antifungal drug responses, we observed functional divergence in filamentation regulation between the two alleles of *ERG251* (*ERG251-A* and *ERG251-B*) with *ERG251-A* exhibiting a dominant role in the SC5314 genetic background. Finally, in a murine model of systemic infection, homozygous deletion of *ERG251* resulted in decreased virulence while the heterozygous deletion mutants maintain their pathogenicity. Overall, this study provides extensive genetic, transcriptional and phenotypic analysis for the effects of *ERG251* on drug susceptibility, fitness, filamentation and stress responses.

**AUTHOR SUMMARY:** Invasive infections caused by the fungal pathogen *Candida albicans* have high mortality rates (20-60%), even with antifungal drug treatment. Numerous mechanisms contributing to drug resistance have been characterized, but treatment failure remains a problem indicating that there are many facets that are not yet understood. The azole class of antifungals targets production of ergosterol, an essential component of fungal cell membranes. Here, we provide insights into the contributions of *ERG251,* a component of the ergosterol biosynthesis pathway, to increased growth in azoles as well as broad scale effects on stress responses filamentation and pathogenicity. One of the most striking results from our study is that even a single nucleotide change in one allele of *ERG251* in diploid *C. albicans* can lead to azole tolerance. Tolerance, a distinct phenotype from resistance, is the ability of fungal cells to grow above the minimum inhibitory concentration in a drug concentration-independent manner. Tolerance frequently goes undetected in the clinic because it is not observable in standard assays. Strikingly, azole tolerance strains lacking one allele of *ERG251* remained virulent in a mouse model of infection highlighting the potential for mutations in *ERG251* to arise and contribute to treatment failure in patients.

## INTRODUCTION

*Candida albicans* is the most prevalent human fungal pathogen, affecting millions of people and causing serious and deadly infections in immunocompromised and immunodeficient individuals [1–4]. Invasive infections caused by *C. albicans* can result in mortality rates nearing ∼60% despite the existing antifungal therapies [1,2,5,6]. Antifungal treatment failures and infection recurrences are common [1,2,7,8]. One contribution to treatment failure is drug resistance, which is defined as the ability to grow above the minimum inhibitory concentration (MIC) of a drug-susceptible isolate at rates similar to growth in the absence of drug. However, treatment failure can also occur in strains that are classified as susceptible based on MIC. This highlights the importance of drug tolerance, which is the ability of a fungus to grow slowly above the MIC in a drug concentration-independent manner [7,9,10].

Fungistatic azole drugs target Erg11 in the ergosterol biosynthesis pathway [6,11–13]. Ergosterol is an essential component of fungal cell membranes and acts to maintain cell membrane integrity and fluidity. Azole exposure leads to the depletion of ergosterol and accumulation of a toxic sterol 14-ɑ-methylergosta-8,24(28)-dien-3β,6α-diol (herein referred to dienol) that permeabilizes the plasma membrane, arrests fungal growth, and increases sensitivity to environmental stresses [12,14,15]. During treatment with fungistatic azoles, many *Candida* species can rapidly evolve drug resistance through various mechanisms including modification or overexpression of the gene encoding the drug target *ERG11* and upregulation of drug efflux pumps encoded by *CDR1*, *CDR2*, and *MDR1* [16,17]. However, these are not the only possible mechanisms. For example, the transcription factor Adr1 has recently been identified as a key regulator of ergosterol biosynthesis and hyperactivation of Adr1 conferred azole resistance in *C. albicans* [18].

Ergosterol biosynthesis is broadly conserved among the Saccharomycotina phylogeny group which includes *Candida* species as well as the baker’s yeast *Saccharomyces cerevisiae.* However, some differences in gene duplication and expression patterns in the more than 20 enzymes along the ergosterol biosynthetic pathway have been identified [19–22]. Ergosterol biosynthesis is divided into three parts: the mevalonate, late, and alternate pathways [21,23]. The mevalonate pathway is responsible for the production of farnesyl diphosphate (FPP), an important ergosterol intermediate. Dephosphorylation of FPP generates farnesol, a quorum-sensing molecule that can regulate the yeast-to-hyphae transition and biofilm formation in *C. albicans* [24–26]. The late pathway is responsible for using FPP to synthesize ergosterol. The rate-limiting enzyme, lanosterol 14-α-demethylase, Erg11 is the direct target of azoles. Inhibition of Erg11 by azoles activates the alternate pathway that proceeds from lanosterol toward the production of the toxic sterol dienol. Key enzymes in the alternate pathway, Erg6 and Erg3, are respectively responsible for the initial step and last step of dienol generation [14,21,27]. Inactivation or modification of *ERG3* or *ERG6* impacts drug susceptibility of many *Candida* species [14,20,22,28–33]. For example, loss of Erg6 function reduces susceptibility to nystatin and polyenes in *C. glabrata* [28,34,35]. Loss of Erg3 function confers resistance to azoles in *C. albicans*, *C. parapsilosis*, *C. dubliniensis* and resistance to polyenes in *C. albicans* and *C. lusitaniae* [22,29–31,36,37]*. ERG3* inactivation causes the depletion of ergosterol and accumulation of 14α-methylfecosterol which supports growth in presence of azoles despite altered membrane composition [14,22].

Both the late and alternate pathways of ergosterol biosynthesis utilize C-4 sterol methyl oxidase, the catalytic component of the C-4 demethylation complex that is responsible for removing the two methyl groups from the C-4 position of the sterol molecule [20,21,38]. In *Saccharomyces cerevisiae,* Erg25 is the solo C-4 sterol methyl oxidase and essential for standard growth [21,39]. However, both *Aspergillus fumigatus* and *C. albicans* encode two membrane C-4 sterol methyl oxidases with one of them serving as the primary enzyme during biosynthesis [19,20,38]. In *C. albicans*, *ERG251* encodes the primary C-4 sterol methyl oxidase, but few studies have characterized it independently and the findings are contradictory. For example, *erg251Δ/Δ* exhibited increased fluconazole (FLC) susceptibility and accumulation of eburicol, the direct precursor for 14α-methylfecosterol, in the presence of FLC [20]. However, in a haploid *C. albicans* strain, transposon insertion into *ERG251* resulted in decreased FLC susceptibility [36]. This contradiction highlights the need to understand the effect of growth conditions and genetic background on the relationship between *ERG251* and drug susceptibility.

Proper cellular ergosterol levels are crucial for the multiple cellular functions including stress response, nutrient transport, and host-pathogen interactions [21,38]. Deletion or overexpression of the key genes in the ergosterol biosynthetic pathway disrupts ergosterol biosynthesis and results in increased susceptibility to osmotic and cell wall stress [21,40]. Disruption of *ERG6* and *ERG24* also leads to the reduced transport of potassium, calcium and metal in *S. cerevisiae* and *A. fumigatus* [41–43]. Furthermore, *C. albicans erg2Δ/Δ* and *erg24Δ/Δ* mutants exhibit abnormal vacuolar physiology and filamentation defects, and are avirulent in disseminated model of candidiasis [44].

In this report, we determined the effects of heterozygous and homozygous inactivation of *ERG251* on drug susceptibility, filamentation, virulence, and response to stress. Heterozygous inactivation of *ERG251* in *C. albicans* resulted in azole tolerance. Strikingly, we identified recurrent heterozygous point mutations in *ERG251* in two distinct genetic backgrounds (SC5314 and P75063) in three independent FLC evolution experiments [10] (Zhou et al. 2024, under review). Evolved strains and engineered strains with *ERG251* point mutations were either tolerant or resistant to azoles. Azole tolerance occurred with single allele dysfunction of *ERG251* in both of the euploid genetic backgrounds. Azole resistance occurred with single allele dysfunction of *ERG251* in combination with concurrent aneuploidy of chromosome 3 and chromosome 6. Homozygous deletion of *ERG251* resulted in increased fitness in the presence of low concentrations of FLC, but decreased fitness in rich medium, especially at low cell density. During FLC exposure, *ERG251* deletion mutants (heterozygous and homozygous) exhibited upregulation of the Zinc finger transporter, *ZRT2,* and genes in the alternate sterol biosynthesis pathway. Importantly, overexpression of *ZRT2* was sufficient to increase azole tolerance, although to a lesser extent than the *ERG251* single allele dysfunction mutants. Therefore, we propose that the combination of upregulated *ZRT2* and the alternate sterol pathway drive *ERG251*-mediated azole tolerance. Interestingly, we found that deletion of only the A allele of *ERG251* (*ERG251-A*), not the B allele, resulted in a filamentation defect in SC5314 background. Lastly, *erg251Δ/Δ* showed decreased virulence while both heterozygous deletion mutants maintained their pathogenicity, highlighting the importance of Erg251 in virulence. In summary, we showed the complex and pleiotropic effects of Erg251 on fitness, stress responses, filamentation, and pathogenicity while also highlighting the importance of Erg251-driven azole drug tolerance.

## RESULTS

### Recurrent point mutations in *ERG251* evolve during adaptation to fluconazole

During *in vitro* evolution of *C. albicans* in the presence of FLC, *ERG251* point mutations were recurrently detected in three independent experiments and within two distinct genetic backgrounds: P75063 and SC5314 (Table 1) [10]. Using whole genome sequencing (WGS), we detected *ERG251* point mutations in 16 FLC-evolved strains (Table 1). In P75063 *ERG251* is homozygous, while in the type strain SC5314, there are two non-synonymous variants between *ERG251-A* and *ERG251-B,* and *de novo* point mutations were identified in both alleles. The point mutations were all heterozygous and were characterized as missense (A60D, A268D, G62W, W265G, and H274Q), nonsense stop gained (L113* and E273*), frameshift (S27fs), or stop lost (*322Y) (Table 1). Many of these *ERG251* point mutations arose in evolved strains with large genomic copy number changes, predominantly whole chromosome trisomies (Table 1 and Fig S1A). The high frequency of mutations in *ERG251* during exposure to antifungal drug suggests an important role of *ERG251* in response to antifungal drug stress.

**Table 1.** List of FLC-evolved strains with different *ERG251* point mutations. *ERG251* mutations were identified in three independent FLC evolution experiments with two distinct genetic backgrounds (SC5314-derived Sn152 and BWP17, and P75063). Multiple single-colony strains from the same evolved population are organized into the same row. *ERG251* point mutations are annotated on the mutated allele, and represented as allele A/allele B, except for AMS5615 where the chromosome region containing *ERG251* became homozygous prior to acquisition of the point mutation. *ERG251* in the P75063 background is homozygous.

| Strains | Genetic background | *ERG251* genotypes | Mutation types | Copy number changes | FLC MIC (24hr) | FLC SMG (48hr) |
| --- | --- | --- | --- | --- | --- | --- |
| **Strains from evolution experiment 1 (Zhou et al. 2024, under review)** |  |  |  |  |  |  |
| Evolved 1.1/1.2 | Sn152 | *ERG251/ERG251<sup>L113\*</sup>* | Stop gained | Chr3x3, Chr6x3, Chr5 LOH | >256µg/ml | N. A |
| Evolved 2.1/2.2 | Sn152 | *ERG251<sup>E273\*</sup>/ERG251* | Stop gained | Chr3x3, Chr6x3 | >256µg/ml | N. A |
| Evolved 3.1/3.3 | Sn152 | *ERG251<sup>A60D</sup>/ERG251* | Missense | Chr3x3, Chr6x3 | >256 µg/ml | N. A |
| Evolved 3.2 | Sn152 | *ERG251/ERG251<sup>\*322Y</sup>* | Stop lost | Chr6x3 | 1 µg/ml | 0.58 |
| **Strains from evolution experiment 2** |  |  |  |  |  |  |
| AMS5615 | BWP17 | *ERG251/ERG251<sup>A268D</sup>* | Missense | Chr4 (partial)x4, Chr7x3, Chr7 LOH | 1µg/ml | 0.79 |
| AMS5617/5618 | BWP17 | *ERG251<sup>S27fs</sup>/ERG251* | Frame shift | Chr4x3,Chr5 LOH | 1µg/ml | 0.73 |
| AMS5622/5623/5624 | BWP17 | *ERG251<sup>G62W</sup>/ERG251* | Missense | Chr7x3 | 1µg/ml | 0.54 |
| AMS5625/5626 | BWP17 | *ERG251<sup>W265G</sup>/ERG251* | Missense | None | 1µg/ml | 0.50 |
| **Strains from evolution experiment 3 [10]** |  |  |  |  |  |  |
| AMS4130 | P75063 | *ERG251/ERG251<sup>H274Q</sup>* | Missense | None | 1µg/ml | 0.26 |

### Single allele dysfunction of *ERG251* leads to azole drug tolerance

To determine the impact of *ERG251* point mutations on drug susceptibility, we quantified azole resistance and tolerance in the evolved isolates. All FLC-evolved strains carrying *ERG251* point mutations in the SC5314-derived background were either resistant (minimal inhibitory concentration, MIC≥ 256µg/ml) or tolerant (Supra-MIC growth, SMG >0.50) to FLC and other azoles (voriconazole, VOC, and itraconazole, ITC) (Table 1, S1 Table). Similarly, the FLC-evolved strain carrying an *ERG251* point mutation in the P75063 background had increased tolerance relative to the progenitor (increase in SMG from 0.13 to 0.26) (Table 1, S1 Table).

Next, we engineered representative point mutations from the FLC-evolved strains into the wildtype, drug-sensitive SC5314 background and determined azole susceptibility. Four different heterozygous *ERG251* point mutations (L113*, W265G, E273*, and *321Y) were selected to represent the range of drug tolerant and resistant phenotypes. The engineered point mutants resulted in a 2-fold increase in MIC and more than an 8-fold increase in tolerance to three different azole drugs (FLC, VOC, ITC) (Fig 1A, S1B and S1 Table).

**Fig 1.**
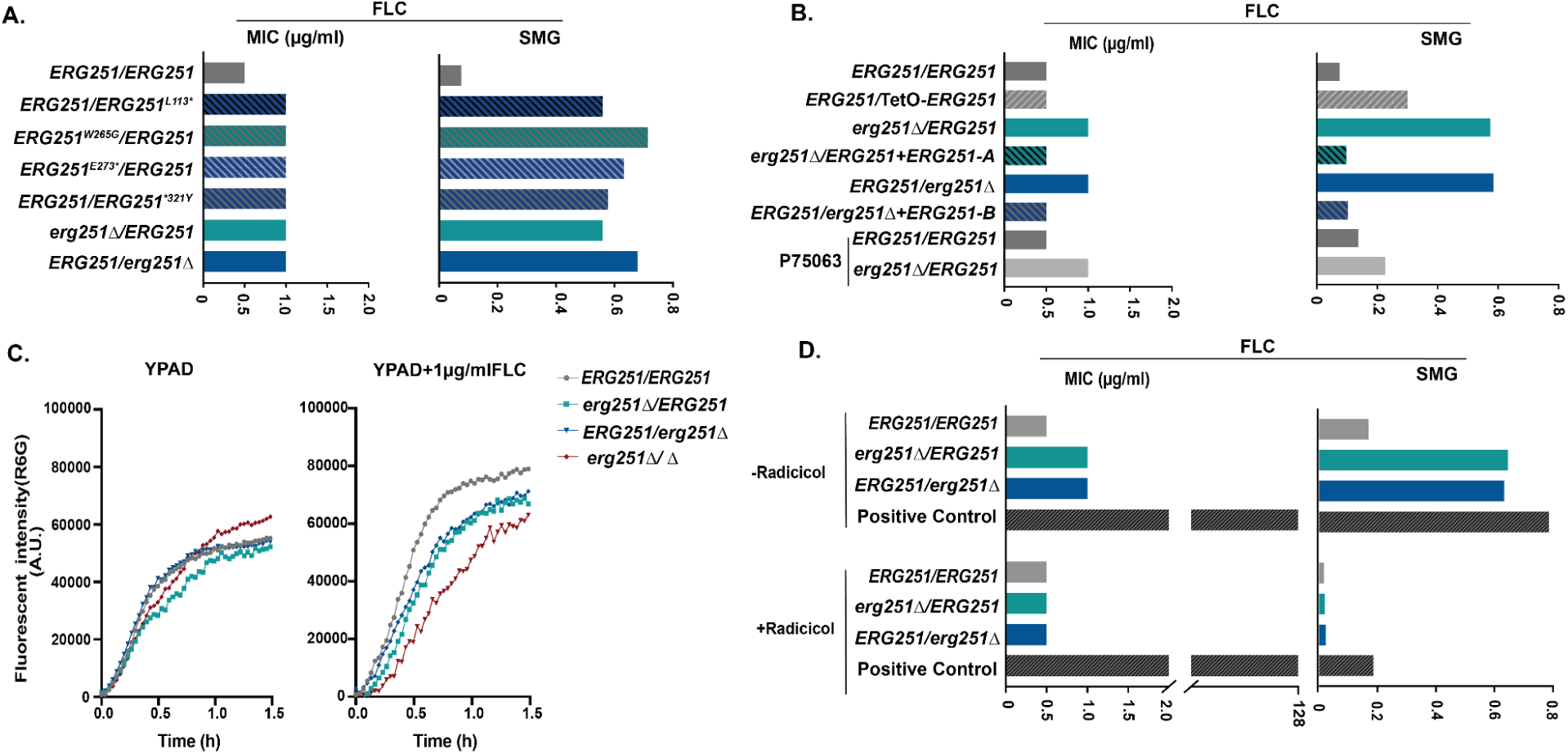
The point mutation of *ERG251* leads to the partial dysfunction of *ERG251* causing acquisition of azole tolerance. Liquid microbroth drug susceptibility assay. Fluconazole (FLC) resistance quantified as the MIC_50_ at 24hr in increasing centrations of FLC (left) and FLC tolerance quantified as the Supra-MIC growth at 48hr (SMG, right) which is the average growth above the MIC_50_ for: **A.** the wildtype SC5314 (*ERG251/ERG251*), engineered heterozygous *ERG251* point mutations strains in the SC5314 background, and both heterozygous deletion mutants of *ERG251* in the SC5314 background; and **B**. the wildtype SC5314 (*ERG251/ERG251*), *ERG251* overexpression strain, both heterozygous deletion mutants of *ERG251* and their corresponding complementation strains, an *ERG251* heterozygous deletion in the P75063 background and wildtype P75063 (P75063-*ERG251/ERG251*) as a control. **C.** Rhodamine 6G efflux kinetics of two heterozygous deletion mutants and the homozygous deletion of *ERG251* in SC5314 with SC5314 (*ERG251/ERG251*) as the control in YPAD (left) and YPAD+1µg/ml FLC (right). Plots indicate average fluorescence intensity changes of Rhodamine 6G (R6G) from three biology replicates over 90 min. **D.** 24hr MIC (left, µg/ml) and 48hr SMG (right, tolerance) in FLC with or without radicicol (Hsp90 inhibitor) treatment for two het deletion mutants of *ERG251* with SC5314 (*ERG251/ERG251*) and a positive control strain known to be resistant to FLC as the controls. **A&B&D**. Each bar represents the average of three technical replicates of a single strain.

Based on these phenotypes we hypothesized that the heterozygous point mutations were due to a loss of gene function. We generated two different heterozygous deletion mutants of *ERG251* by deleting either the A or the B allele in the SC5314 background (a*Δ*/B: *erg251Δ/ERG251* and A/b*Δ*: *ERG251/erg251Δ*). Additionally, we constructed a strain with heterozygous over-expression of *ERG251* (*ERG251*/TetO-*ERG251*) in the SC5314 background. We validated the deletion mutants using WGS and confirmed that transformation did not introduce off-target effects (Fig S1B). Heterozygous deletion of *ERG251* resulted in azole tolerance levels that were the same as all four engineered strains with heterozygous *ERG251* point mutations (Fig 1A and S1 Table). Meanwhile, over-expression of *ERG251* only resulted in a small increase in FLC tolerance (SMG=0.3, Fig 1B). Complementation of the heterozygous mutants *erg251Δ/ERG251* and *ERG251/erg251Δ* with the missing *ERG251* allele (*erg251Δ/ERG251*+*ERG251-A* and *ERG251/erg251Δ*+*ERG251-B*) eliminated the FLC tolerance (Fig 1B and S1B). In the P75063 genetic background, heterozygous deletion of *ERG251* was sufficient to cause the increase in azole tolerance observed for the FLC-evolved strain that carried the *ERG251* point mutation (Fig 1B, Table1). Therefore, we conclude that these *ERG251* point mutations lead to the single allele dysfunction of *ERG251* which causes azole tolerance in *C. albicans*.

We next tested if *ERG251*-driven azole tolerance was caused by upregulated drug efflux pumps and if it is dependent on Hsp90, an important mediator for drug-tolerance and stress response [45,46]. Measurement of Rhodamine 6G (R6G) is a useful method for quantifying efflux pump activity [47]. We found that *ERG251*-driven azole tolerance was independent of drug efflux pumps as indicated by no increase in the rate of efflux of R6G for *ERG251* heterozygous deletion mutants compared to *ERG251/ERG251* (SC5314) during the exposure to FLC (Fig 1C). We found that *ERG251*-mediated tolerance depends on Hsp90 function. Addition of an Hsp90 inhibitor (radicicol, 2.5µM) to assays measuring azole resistance (MIC_50_) and tolerance (SMG) blocked the acquired azole tolerance of *ERG251* heterozygous deletion mutants, but did not inhibit growth of a positive control strain known to be resistant to FLC (Fig 1D).

Additionally, for susceptible and tolerant strains, inhibition of Hsp90 caused FLC to become fungicidal as no viable cells were recovered from higher FLC concentrations combined with radicicol (Fig S2). Hsp90 regulates cell morphogenesis and cell wall stress through the calcineurin pathway, suggesting that *ERG251-*mediated tolerance may also alter cell membrane and/or cell wall stress responses [46].

### Single allele dysfunction of *ERG251* and concurrent aneuploidy leads to azole resistance

The single allele dysfunction of *ERG251* was sufficient to reproduce the azole tolerance phenotype observed in 10/16 of the FLC-evolved strains with *ERG251* point mutations. However, 6 of the FLC-evolved strains with *ERG251* point mutations acquired *bona fide* azole resistance (MIC >256 µg/ml FLC, Table 1). All 6 resistant *ERG251* mutants also had chromosome (Chr)3 and Chr6 concurrent aneuploidies (Fig 2A, Table 1, Evolved 1.1/1.2, 2.1/2.2 and 3.1/3.3 representing two single colonies from three independent FLC-evolved lineages). Recently, we showed that Chr3 and Chr6 concurrent aneuploidy causes azole tolerance and correlates with elevated expression of drug responsive genes located on these chromosomes, including *CDR1*, *CDR2, MDR1*, and *MRR1* (Zhou et al. 2024, under review). We hypothesized that heterozygous deletion of *ERG251* in the Chr3 and Chr6 aneuploid background would make these tolerant cells resistant. To test this hypothesis, we isolated an azole tolerant strain with Chr3 and Chr6 concurrent aneuploidies and wild-type alleles of *ERG251* in the SC5314-derived genetic background (Evolved 4.1: *ERG251/ERG251)* (Fig 2B). We deleted one copy of *ERG251* (on Chr4) from this concurrent aneuploid strain, and confirmed that the mutants maintained the aneuploid chromosomes by whole genome sequencing. Heterozygous deletion of *ERG251* in the concurrent aneuploidy background with elevated drug efflux resulted in a 256-fold increase in MIC, reproducing the azole resistance phenotype observed for the FLC-evolved resistant strains (>256µg/ml, Fig 2B, 2C, 2D and S1 Table). We therefore conclude that the combination of the single allele dysfunction of *ERG251* and concurrent aneuploidy leads to *bona fide* drug resistance.

**Fig 2.**
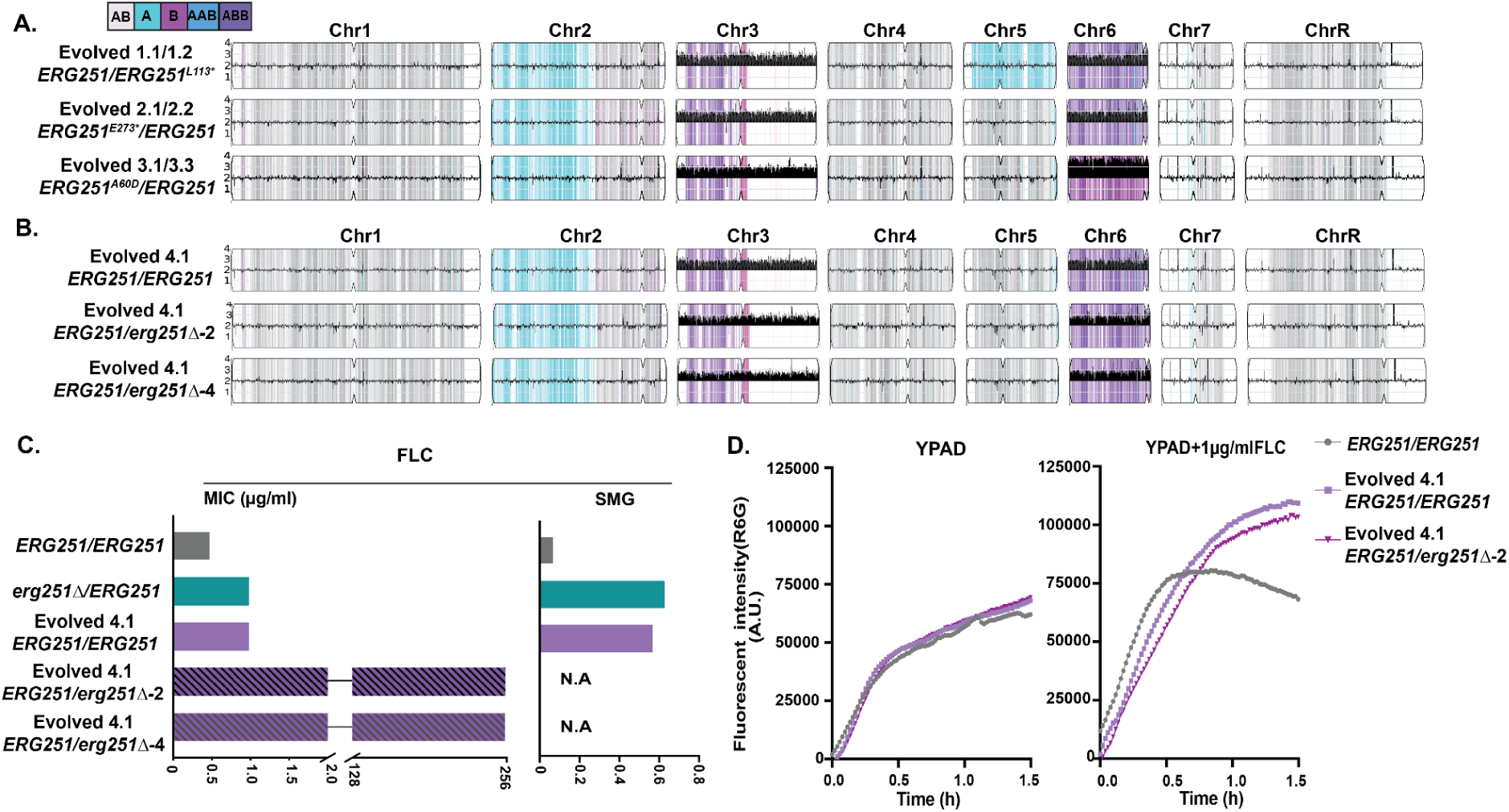
Single allele dysfunction of *ERG251* in combination with concurrent aneuploidy causes azole resistance. A. Representative whole genome sequencing (WGS) data of the FLC-evolved strains 1.1/1.2, 2.1/2.2, and 3.1/3.2 that acquired heterozygous point mutations at *ERG251* and Chr3 and Chr6 concurrent aneuploidy. **B.** WGS data of FLC-evolved strain 4.1 that had wild-type alleles of *ERG251/ERG251* and Chr3 and Chr6 concurrent aneuploidy, plus two *ERG251* heterozygous deletion mutants engineered in the Evolved 4.1 aneuploid background. **A**&**B** WGS data are plotted as the log2 ratio and converted to chromosome copy number (y-axis, 1-4 copies) as a function of chromosome position (x-axis, Chr1-ChrR). The baseline ploidy was determined by propidium iodide staining (S1 Table). Haplotypes relative to the reference genome SC5314 are indicated. **C.** 24hr MIC (left, µg/ml) and 48hr SMG (right, tolerance) in FLC for SC5314 (*ERG251/ERG251)*, *ERG251* heterozygous deletion mutant in the SC5314 background, FLC-evolved strain 4.1, and two *ERG251* heterozygous deletion mutants engineered in the Evolved 4.1 aneuploid background (two independent transformations). Each bar represents the average of three technical replicates per strain. **D.** Rhodamine 6G efflux kinetics of *ERG251* heterozygous deletion mutant in evolved strain 4.1 background with evolved strain 4.1 and SC5314 (*ERG251/ERG251*) as the controls in YPAD (left) and YPAD+1µg/ml FLC (right). Plots indicate fluorescence intensity changes of Rhodamine 6G (R6G) over 90 min.

### Erg251 exhibits contrasting effects on fitness in the presence or absence of drug

Our results show that disruption of one copy of *ERG251* results in tolerance or resistance to azoles in distinct genetic backgrounds. Although we have shown that Hsp90 is required for tolerance, the full range of mechanisms and impact of changing *ERG251* are not known. In order to fully understand the function of *ERG251* in different cellular processes, two independent homozygous *ERG251* deletion mutants were generated in SC5314 background. We confirmed that these deletions did not introduce any large-scale genomic changes (LOH and aneuploidy) (Fig S1B), and two independent *erg251Δ/Δ* mutants (d51 and d70) exhibited identical phenotypes. Both e*rg251Δ/Δ* mutants had decreased growth in rich medium with low initial cell density (OD_600_=0.001) compared to the wild-type control in a 96-well plate format with shaking (Fig 3A and B). Importantly, we found that decreased growth of *erg251Δ/Δ* in YPAD was related to the cell density. The growth defects of *erg251Δ/Δ* were partially rescued by simply increasing initial cell density (from OD_600_=0.001 to 0.005 or 0.01), which mostly restored carrying capacity but not doubling time (Fig 3A and B). In *C. albicans*, cell density is communicated and linked with gene expression via the quorum sensing process, and farnesol is a major quorum sensing molecule secreted by *C. albicans* [24,48,49]. The production of farnesol requires the dephosphorylation of FPP, the precursor for the ergosterol biosynthesis pathway [25,50]. Therefore, we tested the impact of different concentrations of farnesol (0-1000µM) on the growth of *erg251Δ/Δ* mutants with low initial cell density (Fig 3C, Y-axis). We found moderate concentrations of farnesol (62.5-250 µM) improved the growth of *erg251Δ/Δ* in YPAD, while farnesol had no impact on growth of wild-type control (*ERG251/ERG251*) (Fig 3D, Y-axis). Therefore, homozygous deletion of *ERG251* may result in disrupted ergosterol biosynthesis which subsequently provides negative feedback on farnesol production contributing to the growth defect of *erg251Δ/Δ*.

**Fig 3.**
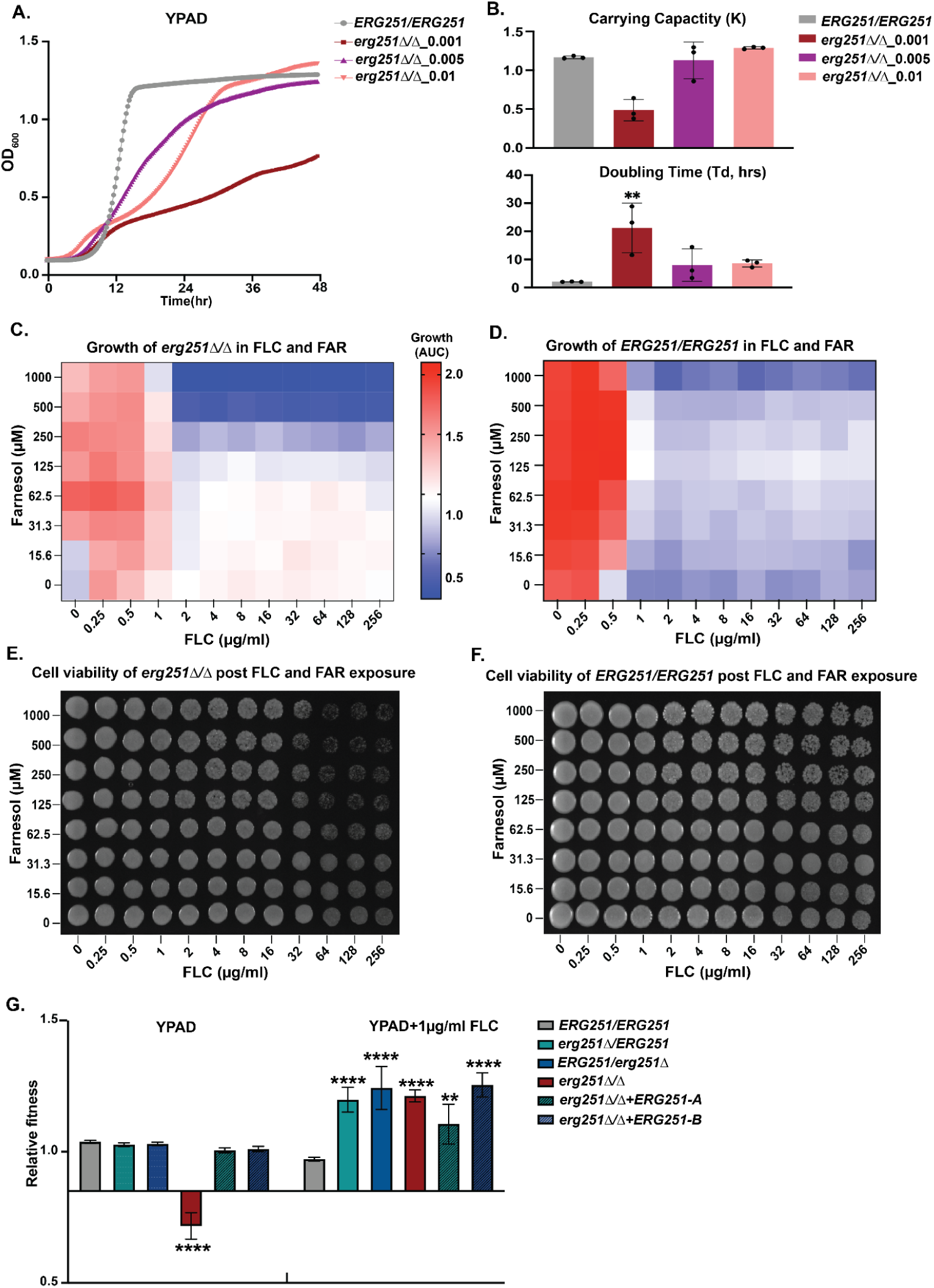
Homozygous deletion of *ERG251* results in decreased fitness at low initial cell density and increased fitness in the presence of low concentrations of FLC (≤1µg/ml). A. 48hr growth curve analysis of *erg251Δ/Δ* started at three different initial cell densities (OD_600_=0.001, 0.005, or 0.01) with *ERG251/ERG251* (SC5314, OD_600_=0.001) as the control. **B.** Carrying capacity (K) and doubling time (Td, hrs) determined from growth curve analysis in Fig 2A. **C&D.** X-Y growth curve assay of (C) *erg251Δ/Δ* and (D) *ERG251/ERG251* in the presence of increasing concentrations of FLC (X-axis, 0-256 µg/ml, 2-fold dilutions) and/or increasing concentrations of farnesol (FAR) (Y-axis, 0-1000 µM, 2-fold dilutions). Growth was estimated with the area under the curve (AUC heatmap) of the 48hr growth curve. **E&F.** Cell viability of (E) *erg251Δ/Δ* and (F) *ERG251/ERG251* after 48 hr exposure to FLC or/and FAR. Cells from Fig 3B were plated on YPAD agar and imaged after 24hr incubation. **G.** Relative fitness calculated from head-to-head competitive assay for *erg251Δ/ERG251, ERG251/erg251Δ, erg251Δ/Δ, erg251Δ/Δ*+*ERG251-A,* and *erg251Δ/Δ*+*ERG251-B* compared to the fluorescent control strain (*ERG251/ERG251*). **B&G**: Values are mean ± SEM calculated from three technical replicates. Data were assessed for normality by Shapiro-Wilk, and significant differences between the *ERG251/ERG251* and mutants were calculated using two-way ANOVA with Dunnett’s multiple comparisons test. \*\*\*\**p*<0.0001, \*\**p*<0.01. **A-G:** At least three biological replicates were performed.

We next measured the impact of FLC on *erg251Δ/Δ* strains. The growth defect of the *erg251Δ/Δ* strain prevented us from conducting MIC and SMG assays for resistance and tolerance because these assays are normalized to growth in rich media (no drug) and *erg251Δ/Δ* strains grow poorly in these conditions. Therefore, we tested the impact of FLC on *erg251Δ/Δ* using a growth curve assay. By monitoring the growth of *erg251Δ/Δ* at different concentrations (0-256µg/ml) of FLC, we found that low concentrations of FLC (≤1µg/ml) increased growth of *erg251Δ/Δ* compared to no drug (Fig 3C, X-axis). High concentrations of FLC (>1µg/ml) had minimal impact on *erg251Δ/Δ* growth (Fig 3C, X-axis). In contrast, the wild-type control (*ERG251/ERG251*) exhibited decreased growth at concentrations at and above the MIC_50_ (0.5µg/ml) (Fig 3D, X-axis). This suggests that total dysfunction of *ERG251* can promote *C. albicans* growth in FLC but only at low concentrations.

Interestingly, adding either farnesol or FLC only partially restored the growth defect of *erg251Δ/Δ* (Fig 3C). Next, we determined if adding farnesol in combination with FLC could further contribute to restored growth of *erg251Δ/Δ*. Growth of *erg251Δ/Δ* from all different concentration combinations showed that low concentrations of FLC (≤1µg/ml) are sufficient to confer increased growth regardless of the concentration of farnesol (Fig 3C). In contrast, high-concentrations of both farnesol (>125µM) and FLC (>1µg/ml) greatly inhibited growth of *erg251Δ/Δ* (Fig 3C). Growth inhibition for the wild-type control (*ERG251/ERG215*) was solely controlled by the FLC concentration (Fig 3D). Furthermore, high-concentration farnesol (>125µM) combined with high concentrations of FLC (>64µg/ml) exhibited a killing effect on the *erg251Δ/Δ* cells but not on the wild-type control (*ERG251/ERG251*) (Fig 3E and F). These results suggest that in the absence of Erg251, farnesol can make FLC fungicidal at high concentrations, likely due to more severe inhibition of cell growth and ergosterol production.

Lastly, a head-to-head competition assay validated the fitness trade-off for *erg251Δ/Δ*, with a fitness cost in YPAD and a fitness benefit in the presence of a low concentration of FLC (1µg/ml) (Fig 3G). This fitness trade-off was not seen for the two heterozygous deletion mutants (*erg251Δ/ERG251* or *ERG251*/*erg251Δ*) and was completely rescued by complementation of the homozygous deletion mutant with either *ERG251-A* or *ERG251-B* (*erg251Δ/Δ+ERG251-A* or *erg251Δ/Δ+ERG251-B*) (Fig 3G). Taken together, we propose that in response to low concentration FLC, *erg251Δ/Δ* may upregulate the alternate sterol production pathway to compensate for ergosterol production and support increased growth (see below).

### Pleiotropic effects of Erg251 on cell wall organization, stress response, and biofilm formation

We next explored the mechanisms by which *ERG251* affects fitness, drug susceptibility, and other stress responses. Transcriptional analysis was performed for the SC5314 wildtype (*ERG251/ERG251)*, two *ERG251* heterozygous deletion mutants, and one homozygous deletion mutant using RNAseq in two different log phase conditions: YPAD and YPAD+1µg/ml FLC. We first focused our analysis on the comparison between *erg251Δ/Δ* and *ERG251/ERG251* in YPAD to understand the role of *ERG251* in a broad range of cellular processes (Fig 4A, S2 Table). Differential expression analysis was used to identify genes with a significant change in abundance in *erg251Δ/Δ* cells compared to *ERG251/ERG251* (913 genes, log2 fold change ≥ 1 or ≤-1 and adjusted p-value < 0.05). Gene Ontology (GO) analyses of differentially expressed genes revealed an overrepresentation of genes associated with cell wall organization, biofilm formation, filamentation growth, metabolic processes, and stress response (Fig 4B, S3 Table).

**Fig 4.**
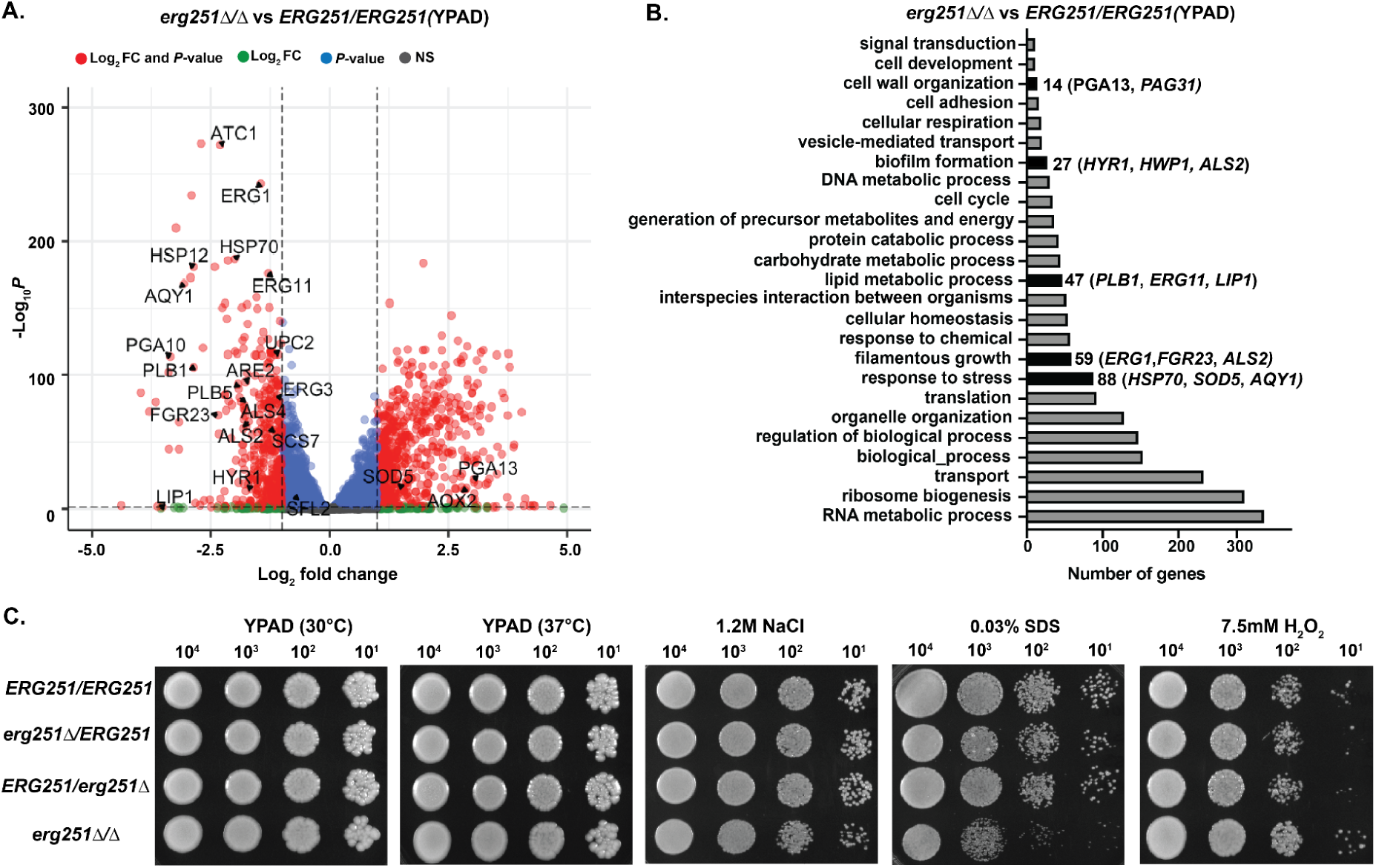
Homozygous deletion of *ERG251* leads to increased sensitivity to cell wall and osmotic stress but decreased sensitivity to oxidative stress. A. Volcano plot for differentially expressed genes (log2 fold change ≥ 1 or ≤-1 and adjusted *p*-value < 0.05) in the *erg251Δ/Δ* mutant compared to *ERG251/ERG251* in YPAD with both fold change and *p*-value indicated. **B**. Gene Ontology (GO) terms for differentially expressed genes (log2 fold change ≥ 1 or ≤ -1 and adjusted *p*-value < 0.05) in the *erg251Δ/Δ* mutant compared to *ERG251/ERG251* in YPAD. **C.** Spot plates growth of *ERG251/ERG251, erg251Δ/ERG251, ERG251/erg251Δ,* and *erg251Δ/Δ* on YPAD (30°C), YPAD (37°C), 1.2M NaCl, 0.03% SDS and 7.5mM H_2_O_2_ agar plates. **A-C**: At least three biological replicates were performed.

We identified many down-regulated genes in *erg251Δ/Δ* cells involved in cell membrane, filamentation, and stress response. Genes that regulate cell membrane structure including lipid metabolism, ergosterol and sphingolipid biosynthesis were down-regulated, including *ERG11*, *ERG1*, *LIP1*, *PLB1*, *UPC2, ARE2,* and *SCS7* (Fig 4A and 4B) [36,51]. Genes that regulate filamentation and biofilm formation were also down-regulated, including *FGR23*, *HYR1*, *ALS2*, and *ALS4* (Fig 4A and 4B) [52–54].

Additionally, the osmotic stress related gene *AQY1* and heat stress related genes *HSP70* and *HSP12* were also down-regulated (Fig 3A) [54–56]. In contrast, genes that are involved in oxidative stress response like *SOD5* and *AOX2* were up-regulated (Fig 3A and 3B) [57,58]. These changes in lipid metabolism and stress response may affect metabolism and nutrient availability more broadly including carbon and amino acid metabolism [20,59].

GO analysis of biological process identified enrichment of 26 genes in *erg251Δ/Δ* cells that encode proteins with GlycosylPhosphatidylInositol (GPI)-anchored motifs. GPI anchors attach proteins to the cell surface contributing to cell-wall integrity, cell-cell interaction, and hyphal formation [60,61]. These genes include *HYR1*, *FGR23*, and *SOD5* as well as cell wall specific genes in the *PGA* and *ALS* families. Overall, genes encoding GPI-anchored motifs were down-regulated in *erg251Δ/Δ* cells (20 out of 26) (Fig 4A and S4 Table) consistent with prior work demonstrating cross-talk between the ergosterol and GPI biosynthesis pathways [62,63].

Phenotypic analysis was consistent with transcriptional analysis for the pleiotropic effects of *ERG251* on cell wall organization and stress responses. Compared to wildtype (*ERG251/ERG251*), *erg251Δ/Δ* exhibited no change in response to increased temperature (37°C). However, *erg251Δ/Δ* exhibited decreased growth in the presence of osmotic (1.2M NaCl) and cell membrane (0.03% SDS) stress, but increased resistance to H_2_O_2_ (Fig 4C). These changes in stress response were not observed for the two *ERG251* heterozygous deletion mutants at either the transcriptional or phenotypic levels (Fig 4C, Fig S3A, and S3B). Taken together, this indicates that the total disruption of *ERG251* results in a dramatic physiological response that impacts cell membrane/cell wall compositions, and osmotic/oxidative stress responses.

### *ERG251-A* exhibits dominant regulation of filamentation

Given that genes related to filamentation and biofilm formation were downregulated in *erg251Δ/Δ*, we next quantified filamentation in all three deletion mutants (homozygous and two heterozygous). Using an *in vitro* filamentation assay, we found that *erg251Δ/ERG251* had a ∼25% decrease in the proportion of hyphae, while *ERG251/erg251Δ* exhibited almost no change compared to wild-type *ERG251/ERG251* (Fig 5A and 5B). Complementation of *erg251Δ/ERG251* with the *ERG251-A* allele restored wild-type filamentation (Fig 5A and 5B). This indicates that *ERG251-A* plays a dominant role in regulating filamentation, while *ERG251-B* is not required for filamentation in *C. albicans*. Additionally, a more severe filamentation defect (∼50%) was observed in *erg251Δ/Δ* compared to *ERG251/ERG251* (Fig 5A and 5B). Complementation of *erg251Δ/Δ* with the *ERG251-A* allele, not the *ERG251-B* allele, was able to partially restore filamentation (Fig 5A and 5B). This data supports a dominant role of *ERG251-A* in regulating filamentation.

**Fig 5.**
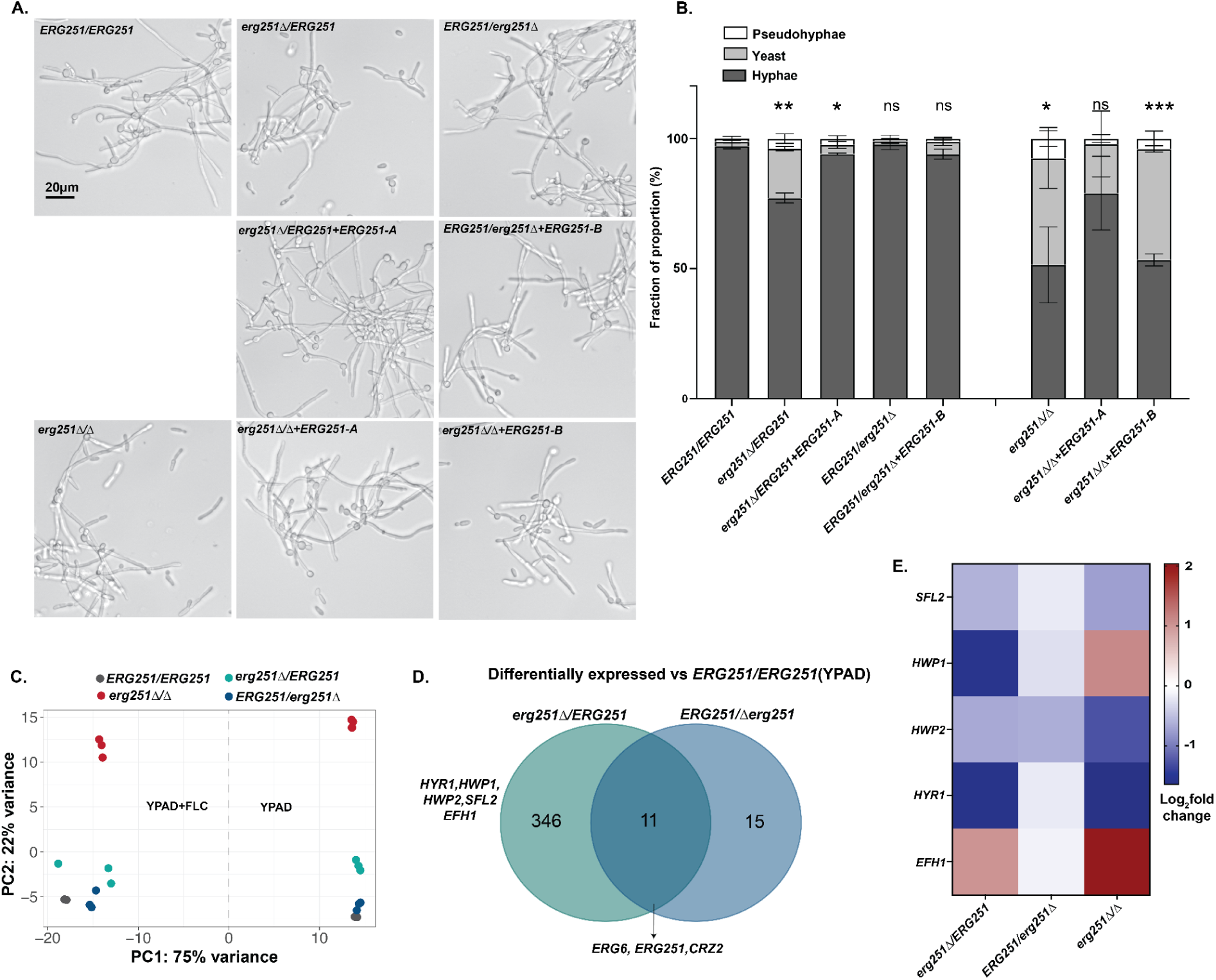
Deletion of ERG251-A but not ERG251-B leads to decreased filamentation. A. Representative filamentation images of wildtype *ERG251/ERG251*, *erg251Δ**/**ERG251, erg251Δ**/**ERG251+ERG251-A, ERG251/erg251Δ, ERG251/erg251Δ+ERG251-B, erg251Δ/Δ, erg251Δ/Δ+ERG251-A,* and *erg251Δ/Δ+ERG251-B*. Cells were induced in RPMI supplemented with 10% FBS for 4 hrs. Scale bar, 20 μm. **B.** Quantification of the yeast (<6μm), pseudohyphae (15-36 μm), and hyphae (>36 μm) from genotypes in Fig 5A. 150 to 500 cells were counted for each strain, and at least two biological replicates were performed. Error bars indicate SEM. Statistical significance for filamentation was compared to *ERG251/ERG251* and assessed using two-way ANOVA with uncorrected Fisher’s LSD, \*\*\**P* <0.001, \*\**P* <0.01,* P ≤ 0.05, ns: *P* >0.05. **C.** Principal component analysis of transcriptional data in YPAD and YPAD+FLC (1µg/ml) for *ERG251/ERG251, erg251Δ**/**ERG251, ERG251/erg251Δ,* and *erg251Δ/Δ*. **D.** Venn diagrams comparing the genes that are differentially expressed in *erg251Δ**/**ERG251* and *ERG251/erg251Δ* (log2 fold change ≥ 0.5 or ≤-0.5 and adjusted *p*-value < 0.1) relative to *ERG251/ERG251* in YPAD. **E.** The relative expression level (log2 fold change) of genes associated with filamentation in *erg251Δ**/**ERG251, ERG251/erg251Δ,* and *erg251Δ/Δ*compared to *ERG251/ERG251* in YPAD.

*ERG251-A* regulation of filamentation might be caused by the control of genes that are involved in the yeast-to-hyphae transition. Transcriptional analysis revealed that deletion of *ERG251-A* (*erg251Δ/ERG251*) resulted in a greater impact on overall gene expression than deletion of *ERG251-B* (*ERG251/erg251Δ*) (Fig 5C, S3A-D). In the YPAD condition, deletion of *ERG251-A* resulted in 357 differentially expressed genes (log2 fold change ≥ 0.5 or ≤-0.5 and adjusted *p*-value < 0.1, S5 Table) compared to deletion of *ERG251-B* which altered expression of 26 genes (log2 fold change ≥ 0.5 or ≤-0.5 and adjusted *p*-value < 0.1, S6 Table)(Fig 5D). Only 11 genes were significantly differentially expressed in both heterozygous mutants including *ERG6*, *ERG251* and *CRZ2* (Fig 5D). Notably, *ERG6* had increased expression in both heterozygous mutants which may contribute to the activation of the alternate pathway for ergosterol biosynthesis (Fig S3A, S3B, and Fig 5D). This suggests there is redundancy of the *ERG251-A* and *ERG251-B* alleles in ergosterol biosynthesis, which is consistent with the same azole tolerance phenotypes we observed upon loss of function of either allele (Fig 1). Furthermore, in the SC5314 background *ERG251-A* and *ERG251-B* had similar RNA abundance (Fig S3E), and both of the tagged proteins, Erg251-A-GFP and Erg251-B-GFP, localized to the endoplasmic reticulum (ER) in both yeast and hyphal phases (Fig S3F). This indicates that the divergent function of the two *ERG251* alleles is not caused by allelic expression or subcellular translocation. Among the 346 genes that were differentially expressed only in *erg251Δ/ERG251*, GO analysis revealed an enrichment of genes that regulate filamentation (Fig S3C, S7 Table). Genes that positively regulate filamentation, including *HYR1* and *HWP1*, and their up-stream transcription factor *SFL2* were all down-regulated in *erg251Δ/ERG251* (Fig S3A and 5E) [53,54]. Transcription factor *EFH1* was up-regulated in *erg251Δ/ERG251* and its overexpression may lead to pseudohyphal formation (Fig S3A and 5E) [64,65]. Finally, we found that in YPAD, both *erg251Δ/ERG251* and *erg251Δ/Δ* have largely conserved regulation of this subset of genes involved in filamentation: *SFL2, HWP1, HWP2, HYR1, HYR3,* and *EFH1* (Fig 5E).

### Homozygous deletion of *ERG251* leads to downregulation of ergosterol biosynthesis genes and upregulation of alternate sterol biosynthesis genes

Homozygous deletion of *ERG251* results in a diverse set of phenotypic effects that may be directly related to disrupted ergosterol biosynthesis and lipid metabolism genes. To more comprehensively analyze the impact of deleting *ERG251* on ergosterol biosynthesis, we analyzed transcription of genes involved in ergosterol biosynthesis from all three pathways: mevalonate, late and alternate (Fig 6A). In YPAD, *erg251Δ/Δ* had decreased expression relative to wild-type for 11 *ERG* genes, and increased expression of *ERG12*, *ERG25*, and *ERG6* (Fig 6A and B). Among the 11 down-regulated *ERG* genes, *ERG1* and *ERG11* had the most significant decreases (log2 fold change= -1.5 and -1.3 respectively). These two genes represent two rate-limiting steps in the ergosterol biosynthesis pathway (Veen M et al., 2003). Two additional key genes were down regulated in *erg251Δ/Δ* compared to wildtype: *UPC2*, encoding a transcription factor that activates *ERG* genes, and *ARE2*, encoding a sterol acyltransferase that regulates the storage and decomposition of ergosterol [66–68].

**Fig 6.**
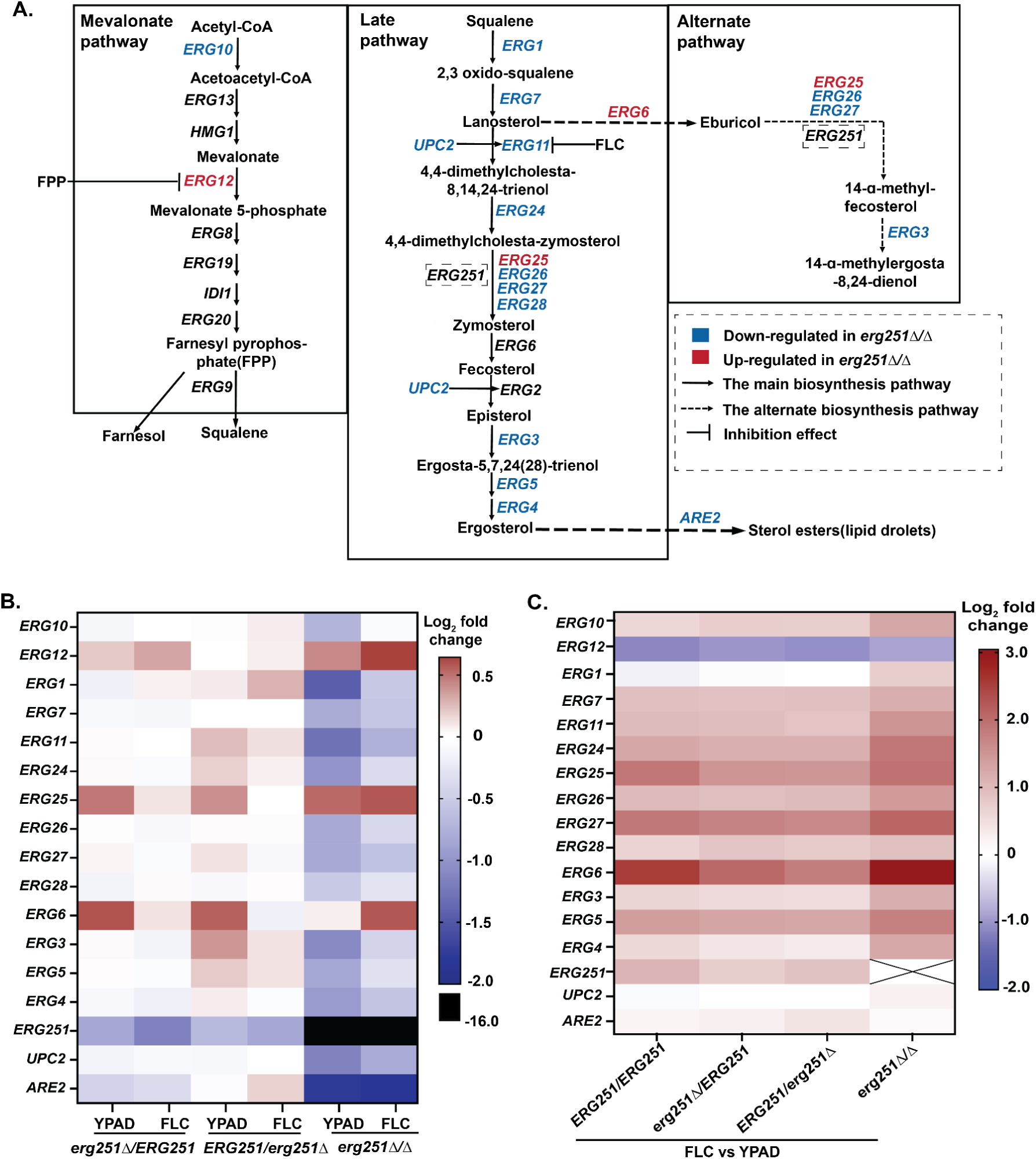
Homozygous deletion of *ERG251* leads to the downregulation of ergosterol biosynthesis genes. A. Overview of the ergosterol biosynthetic pathway in *C. albicans*, including the mevalonate, late ergosterol, and alternate pathways. Genes that were down-regulated (blue) and up-regulated (red) in the *erg251Δ/Δ* under YPAD conditions relative to SC5314 [14,21,69–71]. **B.** The relative gene expression levels (log2-fold change) for all *ERG* genes in the heterozygous and homozygous mutants *erg251Δ**/**ERG251, ERG251/erg251Δ,* and *erg251Δ/Δ* grown in YPAD or YPAD+1µg/ml FLC conditions, compared to the wildtype *ERG251/ERG251* in the same condition. **C.** The relative expression level (log2 fold change) of *ERG* genes in the wildtype *ERG251/ERG251*, and mutants *erg251Δ**/**ERG251, ERG251/erg251Δ,* and *erg251Δ/Δ* grown in YPAD+1µg/ml FLC compared to YPAD condition.

Unlike most ergosterol biosynthesis genes, *ERG25,* the paralog of *ERG251,* had increased expression across both the heterozygous and homozygous *ERG251* deletion mutants (log2 fold change ∼0.5, Fig 6B). *ERG25* is expressed at low levels relative to *ERG251* [20], and increased expression of *ERG25* improved growth and filamentation of the *erg251Δ/Δ* null mutant (Xiong et al, co-submitted manuscript). We therefore hypothesized that Erg25 and Erg251 can compensate for each other during ergosterol biosynthesis despite showing significant sequence divergence (Fig S4A). We were unable to generate a double homozygous deletion of *ERG251* and *ERG25* after multiple attempts. Instead, we generated a heterozygous deletion of *ERG25* in the wild-type and in the *erg251Δ/Δ* mutant, and neither *ERG25* mutant had an effect on drug susceptibility (Fig S4B and S4C). Transcriptional data also supported that the increased abundance of *ERG25* transcripts in *erg251Δ/Δ* was still lower than *ERG251* transcripts in the wild-type background (Fig S4D). Together, our data directly show the paralog compensation between *ERG251* and *ERG25* at the transcriptional level and high-expression level of *ERG251* combined with observed phenotypic changes again supports *ERG251* is the major player for methyl sterol synthesis and can impact drug susceptibility.

When comparing the transcriptional abundance of *ERG* genes between the two growth conditions (FLC vs YPAD), we found that almost all *ERG* genes had increased expression in response to FLC exposure across all four strains, with or without *ERG251* deletion (Fig 6C). Importantly, *ERG* genes that were down-regulated in *erg251Δ/Δ* in YPAD were partially restored with growth in FLC (1µg/ml), which may explain the improved growth of *erg251Δ/Δ* in low concentrations of FLC (≤1µg/ml) (Fig 6C and 3E). Strikingly, *ERG6* had 8-fold increased expression in *erg251Δ/Δ* in the presence of FLC relative to *ERG251/ERG251* (Fig 6C), and over-expression of *ERG6* can result in accumulation of the alternative sterols leading to cell survival in the presence of FLC [21,27,69]. This is consistent with a recent report showing that with FLC exposure, *erg251Δ/Δ* has a depletion of ergosterol and an accumulation of the sterol precursors for the toxic dienol (Lu et al 2023). These results support the homozygous deletion of *ERG251* can lead to upregulation of alternative sterols that promote survival in presence of FLC. However, the heterozygous *ERG251* mutants had minimal differences in *ERG* gene expression (other than *ERG251* itself) compared to the wild-type in the presence of FLC (Fig 6B). This suggests that the azole tolerance of the heterozygous *ERG251* mutants is caused by mechanisms independent from ergosterol biosynthesis gene expression changes.

### Dysfunction of *ERG251* turns on Zinc transporter contributing to decreased azole susceptibility

To determine the mechanism driving decreased drug susceptibility, we further compared the transcriptional analysis of all three *ERG251* deletion mutants during growth in FLC. No significant change in expression was observed for genes encoding the drug efflux pumps *CDR1*, *CDR2*, and *MDR1* across the *ERG251* deletion mutants, compared to wild-type (log2 fold change ≥ 0.5 or ≤-0.5 and adjusted *p*-value < 0.1), with an exception for *MDR1* in *erg251Δ/Δ* that had a 2-fold increase (adjusted *p*-value=9.6×10^-7^) (Fig 7A, S9-S11 Table). We next determined whether there was a conserved transcriptional response across the three *ERG251* mutants after FLC exposure. In YPAD+1ug/ml FLC, only 8 genes were significantly differentially expressed in all three *ERG251* deletion mutants relative to *ERG251/ERG251* (heterozygous deletion mutants: log2 fold change ≥ 0.5 or ≤ -0.5 and adjusted *p*-value < 0.1, homozygous deletion mutant: log2 fold change ≥ 1 or ≤ -1 and adjusted p-value < 0.05) (Fig 5E). Based on the predicted and characterized functions of these 8 genes, we focused on *ZRT2* which encodes a zinc transporter. In *C. albicans*, Zrt2 localizes to the plasma membrane and is essential for Zinc uptake and growth at acidic pH [72]. Interestingly, *ZRT2* was upregulated in both heterozygous and homozygous *ERG251* deletion mutants during FLC exposure. This supports that the dysfunction of *ERG251* can upregulate the Zinc transporter.

**Fig 7.**
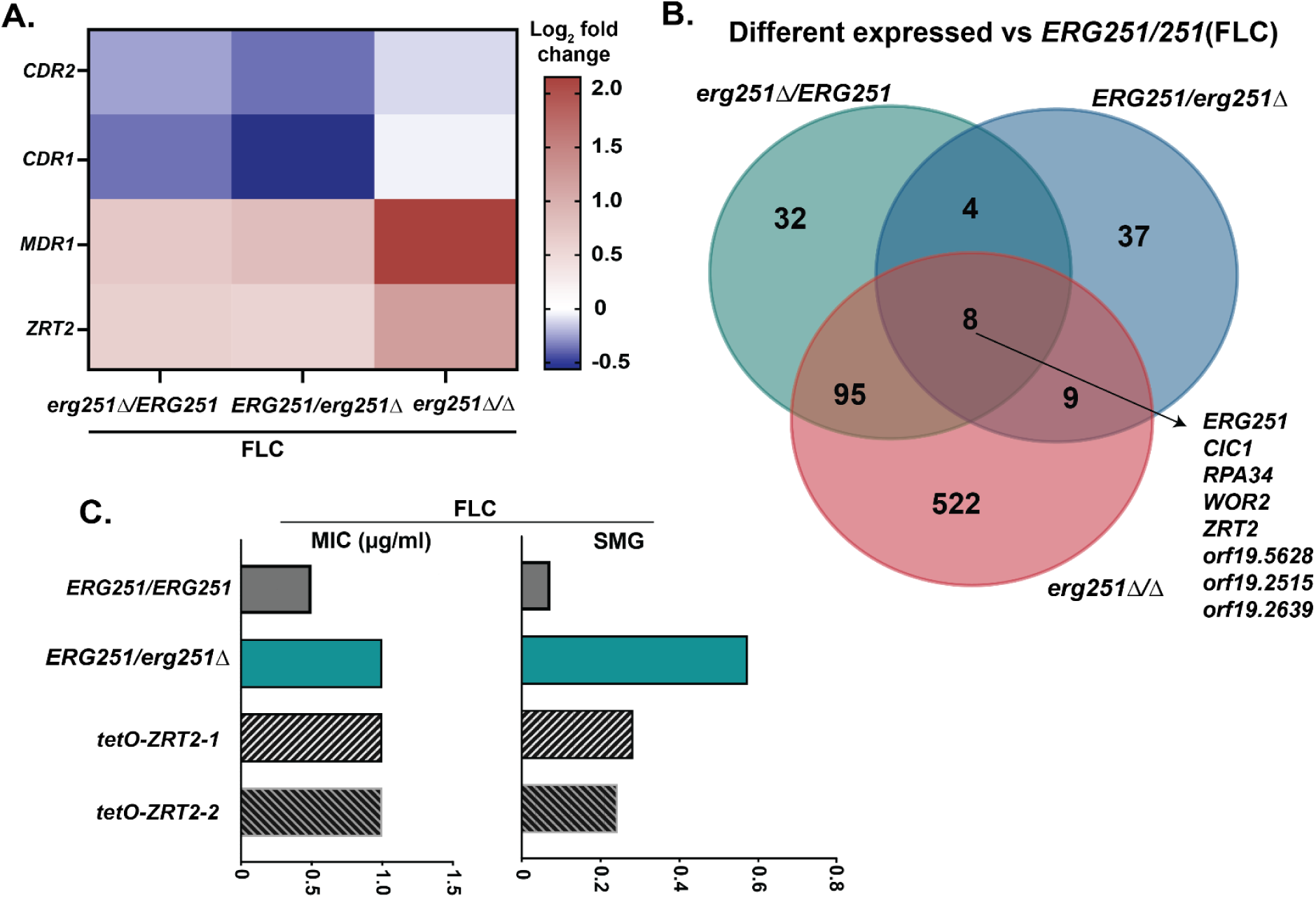
Dysfunction of *ERG251* activates a Zinc transporter contributing to decreased azole susceptibility. A. The relative expression level (log2 fold change) of *CDR1, CDR2, MDR1 and ZRT2* in *erg251Δ**/**ERG251, ERG251/erg251Δ,* and *erg251Δ/Δ* compared to *ERG251/ERG251* under YPAD+1µg/ml FLC condition. **B.** Venn diagrams comparing the genes that differentially expressed in *erg251Δ**/**ERG251, ERG251/erg251Δ* and *erg251Δ/Δ* related to *ERG251/ERG251* under YPAD+1µg/ml FLC condition. **C.** 24hr MIC (left, µg/ml) and 48hr SMG (right, tolerance) in FLC for two *ZRT2* overexpression strains (*tetO-ZRT2*-1 and *tetO-ZRT2*-2) in SC5314 background together with SC5314 (*ERG251/ERG251*) and *erg251Δ**/**ERG251* as the controls. At least three biological replicates were performed.

To determine if elevated *ZRT2* contributes to the azole tolerance in the *ERG251* mutants, we constructed a *ZRT2* overexpression strain. Overexpression of *ZRT2* in the SC5314 background (*ERG251/ERG251*) resulted in 2-fold increase in FLC MIC_50_ and increased FLC tolerance, but resulted in less tolerance than the *erg251Δ/ERG251* mutant (Fig 7C). We propose that Zrt2 may contribute to the drug efflux activities distinct from the ATP-dependent drug efflux pumps such as *CDR1* to increase drug tolerance.

### Single allele dysfunction of *ERG251* maintains pathogenicity in a murine model

We next explored the effects of *ERG251* mutations on pathogenicity to better understand the roles of these mutations during infection. The pleiotropic effects of *ERG251* on varied cellular responses, especially decreased resistance to superoxide and reduced filamentation raise the question of whether dysfunction of *ERG251* would also lead to a pathogenicity defect. We tested the two heterozygous and homozygous *ERG251* deletion mutants in the standard mouse tail-vein injection model of disseminated candidiasis [73]. There was no difference in survival between the wild-type control SC5314 and the heterozygous mutants (e*rg251Δ**/**ERG251* and *ERG251/erg251Δ*) (Fig 8). However, mice infected with *erg251Δ/Δ* had significantly longer survival compared to the wild-type control or either heterozygous deletion mutant with a mean time of survival of 15 days for *erg251Δ/Δ* compared to 8-10 days for all other strains (Fig 8). The statistical analysis showed that the survival curve for *erg251Δ/Δ* is significantly different compared to the wildtype (*P* = 0.0015, Long-rank (Mantel-Cox) test) (Fig 8). We also tested the survival of two complementation strains of *erg251Δ/Δ*, which also had a mean time of survival of 10 days indicating the restoration of virulence (Fig 8). Taken together, this indicates that homozygous deletion of *ERG251* leads to attenuated virulence, which supports the importance of *ERG251* in varied cellular responses essential for pathogenicity. However, the partial dysfunction of *ERG251* has no impact on virulence. The lack of a virulence defect in the heterozygous mutants emphasizes that these mutations, which result in high levels of azole tolerance, remain infectious.

**Fig 8.**
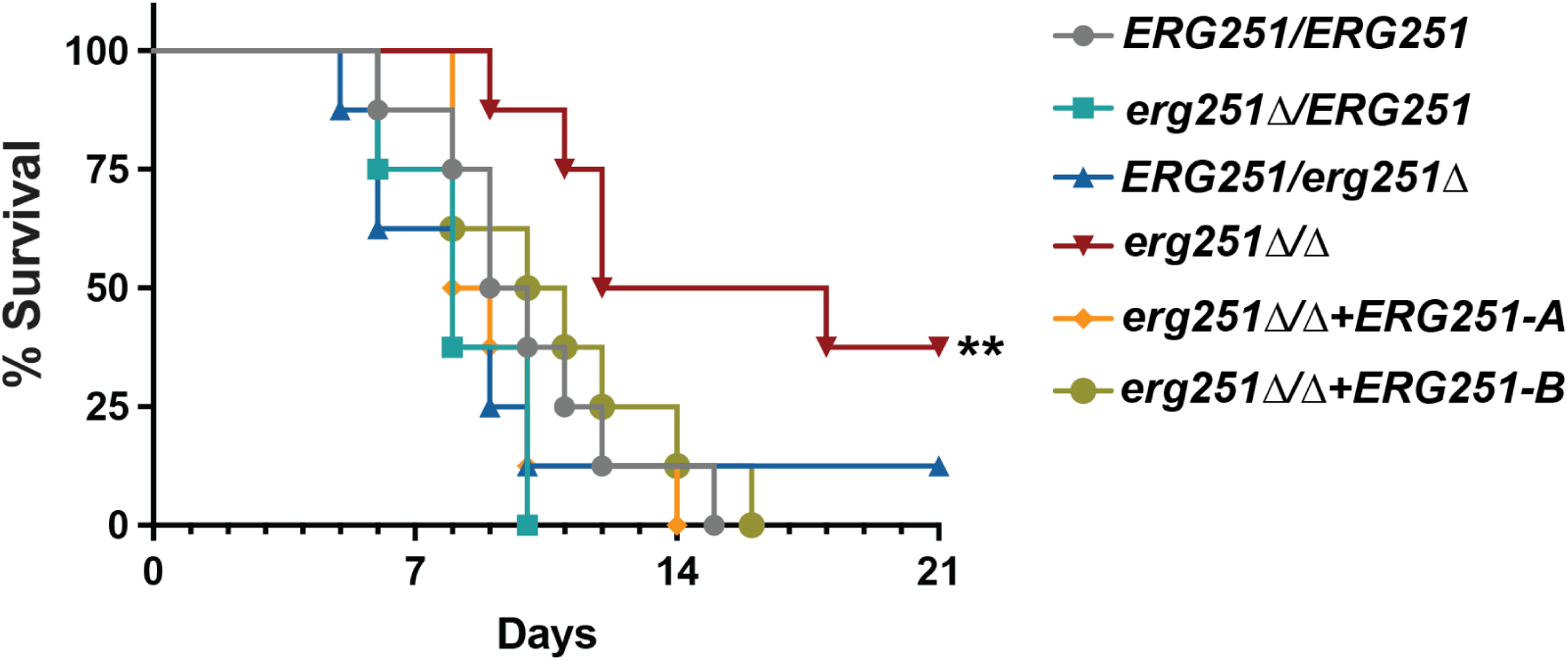
Heterozygous deletion of *ERG251* maintains pathogenicity in a murine model. A. ICR mice were injected via the tail vein with 5×10^5^ cells of *ERG251/ERG251* (SC5314), *erg251Δ**/**ERG251, ERG251/erg251Δ,* and *erg251Δ/Δ*+*ERG251-A* and *erg251Δ/Δ*+*ERG251-B* and survival was presented over the time. The *erg251Δ/Δ* mutant survival curves were significantly attenuated from that of the *ERG251/ERG251* (Log-rank (Mantel-Cox) test; **, *p* = 0.0015). Eight mice per strain were used.

## DISCUSSION

### Changes in *ERG251* impact the susceptibility of *C. albicans* to azoles

Antifungal tolerance varies among *C. albicans* clinical isolates and correlates with the inability to clear an infection. The tolerance phenotype is stable even in the absence of antifungal drug stress [9]. Despite this, the molecular mechanisms causing antifungal tolerance are not known. We found that *ERG251* is a hotspot for point mutations during adaptation to antifungal drug stress, and that heterozygous deletion of *ERG251* can drive azole tolerance in diverse clinical isolates of *C. albicans*. This highlights the novel finding that an *ERG251* point mutation alone can drive azole tolerance and contribute to the adaptation to FLC.

Aneuploidy is associated with the rapid evolution of azole tolerance [74,75] (Zhou et al. 2024, under review). We recently found that the Chr3 and Chr6 concurrent aneuploidies conferred multi-azole tolerance via elevated drug efflux pumps (Zhou et al. 2024, under review). Here, we report that recurrent single nucleotide point mutations in *ERG251* (Chr4) often occur together with aneuploidies of smaller chromosomes (Chr3-Chr7) during adaptation to FLC (Table 1). Therefore, we dissected the impact of aneuploidy and/or *ERG251* point mutations on drug susceptibility by engineering the *ERG251* mutations within euploid or aneuploid backgrounds. Heterozygous point mutations at either allele of *ERG251* in euploid backgrounds led to the single allele dysfunction of *ERG251* which conferred azole tolerance. Heterozygous deletion of *ERG251* in a Chr3 and Chr6 aneuploid background was sufficient to cause very high levels of azole resistance (Fig. 2). In other words, the combination of two mechanisms that independently cause azole tolerance can dramatically increase the growth of fungal cells in drug, resulting in *bona fide* azole resistance. Multiple simultaneous mutations are common in cancer cells and often lead to treatment failure [76–79]. Our results highlight the diverse trajectories that *C. albicans* can take during adaptation to antifungal drugs and support that two independent mechanisms of tolerance (point mutation and aneuploidy) can evolve within the same cell resulting in drug resistance.

The impact of mutations is also influenced by the genetic background of an organism. Across different genetic backgrounds, *ERG251* had conserved roles in causing azole tolerance and driving adaptation to FLC; however, variation in tolerance levels was observed. We identified point mutations in *ERG251* within two distinct genetic backgrounds (SC5314 and P75063) from three independent evolution experiments across multiple labs. All *ERG251* point mutations conferred multi-azole tolerance. However, in the P75063 background, *ERG251*-driven tolerance (SMG=0.26) was lower than in SC5314 background (SMG>0.5). In the P75063 background, *ERG251* is homozygous (no allelic variation) despite the heterozygosity across the rest of Chr4. Additionally, the distinct alleles of *ERG251* in the SC5314 background did not affect azole tolerance (Fig 1A), therefore we hypothesize that genetic variations of *ERG251* between different genetic backgrounds are not causing the differences in tolerance phenotype. Instead, it is more likely to be caused by the genetic variation at other loci in the genome [80].

### Drug susceptibility in heterozygous vs homozygous *ERG251* deletion mutants

One of the particularly striking observations from our study was the range of different heterozygous point mutations that phenocopied the heterozygous deletion of *ERG251*. The numerous possible mutation sites at either allele of the *ERG251* combined with a strong fitness advantage for all characterized *ERG251* mutations in FLC explains why *ERG251* mutations were recurrent in three independent *in vitro* evolution experiments. The probability of a single-nucleotide mutation in *ERG251* that has a phenotype appears to be higher than other types of point mutations that cause increased growth in azoles. For example, point mutations in ergosterol-related genes like *ERG11* and *UPC2* that cause drug resistance in *Candida* species are all homozygous in diploid organisms or the only allele present in haploid organisms [81–85]. Comparatively, all of the *de novo ERG251* point mutations identified here were heterozygous. Although the homozygous deletion of *ERG251* led to a similar fitness advantage as the heterozygous deletion mutants in the presence of low concentrations of FLC (<1µg/ml), the homozygous deletion strain exhibited a fitness cost in rich medium and at higher concentrations of FLC, supporting why only heterozygous mutants were identified during *in vitro* evolution experiments.

Prior studies of the relationship of *ERG251* to azole susceptibility in *C. albicans* have found conflicting results. In some experiments, loss of *ERG251* through genetic manipulation or pharmacological inhibition resulted in increased susceptibility to azoles whereas in other experiments, disruption of *ERG251* decreased susceptibility to azoles [20,36]. Our findings help to explain these disparate results. We observed substantial phenotypic differences between heterozygous and homozygous deletion mutants indicating that the amount of Erg251 remaining in the cell can affect ergosterol biosynthesis and other metabolic processes. Another group recently demonstrated that *erg251Δ/Δ* phenotypes are dependent on other growth conditions, including oxygen availability (Xiong et al. 2024, co-submitted manuscript). We also found that the growth of *erg251Δ/Δ* was dependent on cell density and azole concentration. The *erg251Δ/Δ* mutants had decreased susceptibility to low concentrations of FLC, but increased susceptibility to high concentrations of FLC, especially in combination with the quorum sensing molecule farnesol.

### Partial differentiation of *ERG251-A vs ERG251-B* function

In the SC5314 background, the A and B alleles of *ERG251* differ at two non-synonymous SNPs. When comparing the heterozygous mutations from the evolution experiments and engineered strains, we did not observe any differences between the alleles in growth and azole-related phenotypes. GFP-tagging of both alleles showed similar localization patterns, and RNAseq analysis showed that both alleles are expressed at similar levels in rich media and FLC. However, we observed striking differences between the two alleles when considering overall gene expression patterns and filamentation (Fig. 5). In the heterozygous background (SC5314), *ERG251* exhibited allelic differences in controlling filamentation with *ERG251-A* as the dominant player. Phenotypic analysis showed that deletion of *ERG251-A* alone can lead to filamentation defects while deletion of *ERG251-B* had no effect on filamentation. The filamentation defect was correlated with altered gene expression of filamentation-related genes like *HYR1* and *EFH1* [53,54,64,65]. *ERG251* is another rare example of a gene containing a gain-of-function allele that regulates filamentation in *C. albicans*, similar to the transcription factor Rob1 [86]. However, future work is needed to understand the molecular basis of the functional distinctions between diverse *ERG251* alleles.

### Sterol composition changes when *ERG251* is inactivated

Homozygous deletion of *ERG251* had a global impact on the expression of sterol biosynthetic genes resulting in down-regulation of ergosterol biosynthetic genes and upregulation of alternate sterol biosynthetic genes. Together with data from a previous study and the co-submitted manuscript by Xiong et al., we propose that homozygous deletion can lead to sterol composition changes that include reduced ergosterol and an accumulation of alternative sterols [20](Xiong et al. 2024). These changes in sterol composition are further supported by additional phenotypic changes we observed, including increased cell wall sensitivity of *erg251Δ/Δ*.

We also observed a connection between farnesol and ergosterol biosynthesis. We found moderate concentrations of farnesol (62.5-250 µM) improved the growth of *erg251Δ/Δ* in rich media, while farnesol had no impact on growth of wild-type control. Therefore, homozygous deletion of *ERG251* may result in disrupted ergosterol biosynthesis which subsequently provides negative feedback on farnesol production contributing to the growth defect of *erg251Δ/Δ*. One *ERG* gene, *ERG12,* had decreased expression in response to fluconazole - the opposite trend from all other *ERG* genes (Fig 6). *ERG12* encodes mevalonate kinase and converts mevalonate into 5-phosphomevalonate, the precursor for farnesyl pyrophosphate (FPP). One hypothesis is that the FLC-induced repression of *ERG12* is caused by the inhibition of the farnesol or FPP via negative feedback. We also found that homozygous deletion of *ERG251* caused increased expression of *ERG12*. The increased expression of *ERG12* combined with farnesol-related growth phenotypes in the *erg251Δ/Δ* mutant suggests that *ERG12* expression is negatively regulated by farnesol or its precursor, FPP, and that both deletion of *ERG251* and FLC exposure impact *ERG12* expression possibly via farnesol production.

When *ERG251* was inactivated, its paralog *ERG25* was upregulated to compensate for the loss of the primary enzyme. The shared enzymatic function and regulatory networks of paralogs supports a model where compensation between *ERG251* and *ERG25* occurs on the level of gene expression [38,87]. However, we found that the upregulated *ERG25* didn’t reverse the phenotypic changes observed for *ERG251* deletion mutants because *ERG25* had much lower expression (Fig S4D). Other studies also showed ectopic over-expression of *ERG25* only partially recovered the phenotype changes of *erg251Δ/Δ* [20](Xiong et al. 2024). In conclusion, in *C. albicans* both Erg251 and Erg25 function as the essential C-4 sterol methyl oxidase with Erg251 serving as the primary role controlling drug susceptibility, filamentation, biofilm formation and other stress responses [20](Xiong et al. 2024).

### Upregulated Zinc transporter contributing azole tolerance

We also found transcriptional upregulation of the Zinc transporter, Zrt2, in all *ERG251* deletion mutants (heterozygous and homozygous) during exposure to fluconazole. The upregulation of this plasma membrane transporter could be caused by alterations in cell membrane structure that coincide with the down-regulation of ergosterol biosynthesis in *ERG251* deletion mutants [41–43]. Compared to *ERG251* heterozygous deletion mutants, over-expression of *ZRT2* led to the same 2-fold increase in MIC and a significant, but smaller increase in azole tolerance (Fig. 7). Therefore, we propose that *ERG251*-mediated changes in sterol metabolism and upregulated Zinc transport together promote decreased drug susceptibility observed for the *ERG251* deletion mutants.

In summary, this study identified recurrent heterozygous point mutations in the methyl sterol oxidase *ERG251* during adaptation to antifungal drug stress and characterized for the first time point mutation-driven azole tolerance. We utilized genetic, transcriptional and phenotypic analyses to understand the effects of inactivating *ERG251* both partially and completely. Increased azole tolerance was observed in two distinct genetic backgrounds, and heterozygous mutations of *ERG251* promote multi-azole tolerance while maintaining virulence in a mouse model of systemic infection. Future experiments will explore *ERG251* genetic diversity and phenotypes across a broad range of clinical isolates in *Candida albicans* and in related *Candida* species.

## MATERIALS AND METHODS

### Yeast isolates and culture conditions

All strains used in this study are listed in S1 Table including FLC evolved isolates and engineered yeast and bacteria strains. Strains were stored at -80℃ in 20% glycerol. Isolates were grown in YPAD media (20 g/L peptone, 10 g/L yeast extract, 2% dextrose, and 15 g/L agar for plates) supplemented with 40 µg/ml adenine and 80 µg/ml uridine.

### Strain construction

All engineered strains in this study were generated in the SC5314 background, except one *ERG251* heterozygous deletion in the P75063 background. Strains were constructed by lithium acetate transformation using PCR products with at least 140 bp of homology to the target locus. Primers used in this study are listed in S12 Table.

#### *(i) ERG251* heterozygous deletion

The FLIP-NAT construct was PCR amplified from the plasmid pJK863 [88] using primer sets 1630+1631 and transformed into background strains SC5314 and P75063. NAT-resistant transformants were PCR screened for correct integration of the FLIP-NAT construct at the *ERG251* locus using primer pairs 1652+1045 (left of integration) and 1636+1653 (right of integration). Transformants were validated by whole genome sequencing for correct integration.

#### *(ii) ERG251* homozygous deletion

To promote FLIP-mediated excision of FLIP-NAT, correct heterozygous deletion strains *erg251Δ/ERG251* and *ERG251/erg251Δ* were inoculated in YNB+BSA from frozen stocks and incubated at 30℃, 220rpm, for 48 hrs. Cultures were diluted and 100 cells were plated on YPAD agar, then incubated at 30°C for 24 hrs. Recovered colonies were patched to both YPAD and YPAD+150 µg/ml NAT. Colonies growing on only YPAD were screened for correct FLIP-mediated excision of FLIP-NAT using primer pairs 1574+1575 (inside NAT) and 1652+1653 (across *ERG251*). Colonies that correctly excised FLIP-NAT were re-transformed with the FLIP-NAT construct (PCR amplified from the plasmid pJK863 [88] using primer sets 1630+1631). NAT-resistant transformants were PCR screened for correct integration of the FLIP-NAT construct at the remaining *ERG251* locus using primer pairs 1652+1045 (left of integration), 1636+1653 (right of integration), and 1632+1633 (inside *ERG251*). Transformants were validated by whole genome sequencing for correct integration.

#### *(iii)* Construct *ERG251-NAT* plasmid

To generate *ERG251* mutant complementation and point mutation, we built up an *ERG251-NAT* plasmid by fusing *ERG251* upstream plus gene (1644+1645), *NAT* (1574+1575), and *ERG251* downstream (1646+1647) into the pUC19 backbone (1578+1579). PCR amplified fragments were aligned using NEBuilder HiFi DNA Assembly Cloning Kit following the manufacturer’s instructions and transferred into *E. coli*. Ampicillin-resistant transformants were screened using primer pairs 1352+1353 and saved in frozen stocks as pAS3118.

#### *(iv) ERG251* mutant complementation

The wild-type *ERG251* upstream region and genes (A or B) were PCR amplified from heterozygous deletion strains using primer pair 1652+1645. The *NAT* gene and downstream *ERG251* region were PCR amplified from pAS3118 using 1574+1653 primers. SOEing PCR was performed using primer pair 1652+1653. The subsequent *ERG251-NAT* construct was transformed into the *erg251Δ/ERG251*, *ERG251/erg251Δ,* and *erg251Δ/Δ* mutants that had previously excised *FLIP-NAT* as described above. NAT-resistant transformants were PCR screened for correct integration of the *ERG251-NAT* construct using primer pairs 1634+1154 (left integration), 1636+1635 (right integration), and 1634+1635 (across integration). Transformants were validated by whole genome sequencing for correct integration.

#### *(v) ERG251* point mutation

Site-directed mutagenesis using double-primer PCR was used to generate *ERG251* point mutation construct. Primers with the desired mutations were paired with *ERG251-NAT* upstream (1652) or downstream (1653) primer to amplify mutated *ERG251-NAT* construct from pAS3118.

Four different point mutations were engineered in this study: L113*(1652+1649/1648+1653), W265G(1652+1655/1654+1653), E273*(1652+1651/1650+1653), and *321Y(1652+1657/1656+1653). The amplified two fragments were fused using SOEing PCR and transformed into the SC5314 background. Transformants were first PCR screened using primers 1652+1575 (left integration) and 1636+1653 (right integration) for correct integration and then validated by whole genome sequencing for base substitution and mutated allele.

#### *(vi) ERG251* overexpression

The TetO promoter replacement construct was PCR amplified using primer pair 1679+1680 from plasmid pLC605 [89] and transformed into the SC5314 background strain. NAT-resistant transformants were PCR screened for correct integration of the TetO promoter replacement using primer pairs 1652+1176 (left integration) and 1177+1633 (right integration).

#### (vii) ERG251-GFP

The c-terminal GFP-NAT construct was PCR amplified from plasmid pMG2120 [90] using the 1925+1926 primer pairs and transformed into the SC5314 background strain . NAT-resistant colonies were PCR screened for correct integration of the GFP-NAT construct at the c-terminal end of the *ERG251* locus using primer pairs 1632+1927 (left integration) and 1636+1653 (right integration). Transformants were validated by Sanger sequencing for tagged alleles.

#### *(viii) ERG25* heterozygous deletion

The FLIP-NAT construct was PCR amplified from plasmid pJK863 using primer pairs 1921+1922 and transformed into the SC5314 background strain. NAT-resistant transformants were PCR screened for correct integration of FLIP-NAT at the *ERG25* locus using primer pairs 1923+1045 (left integration) and 1636+1924 (right integration).

##### Filamentation

Strains were inoculated in 2% dextrose YPAD from frozen stocks and incubated at 30℃, 220 rpm for 16 hrs. Strains were diluted 1:100 into RPMI+10% FBS, then incubated at 37℃ for 4 hrs. Cells were harvested, washed once with PBS, and resuspended in PBS before microscopy. Images were captured using an Olympus IX83 microscope and analyzed using ImageJ v1.54d.

##### Microscopy

Erg251-GFP tagged strains were struck on YPAD agar plates from frozen stocks and incubated at 30℃ for 24 hrs. Cultures were inoculated in 2% YPAD and incubated at 30℃, 220 rpm for 16 hrs. Cultures were diluted 1:100 in fresh 2% YPAD or RPMI+10% FBS, then incubated at 30℃, 220 rpm for 4 hrs. Cells were spun down, washed once with PBS, and resuspended in PBS before microscopy. Images were captured using an Olympus IX83 microscope.

##### Spot plate assay

Strains were inoculated in 2% dextrose YPAD from glycerol stocks and incubated at 30℃, 220 rpm for 16 hrs. Cultures were normalized to 10^6^ cells/ml, then 10-fold serially diluted. 10µl of each (10^6^-10^3^) dilutions were spotted onto YPAD agar with and without drugs. All spot plates were performed in triplicates. Plates were incubated for 48 hrs at 30℃ and imaged using a BioRad GelDoc XR+ imaging system.

##### RNA sequencing

i) RNA extraction: For RNA extraction, all 4 strains (wildtype and 3 *ERG251* deletion mutants) were struck on YPAD agar plates from frozen stocks and incubated at 30℃ for 24 hrs. Cultures were then inoculated in 2% YPAD (50 ml) and incubated at 30℃, 220 rpm for 16 hrs. Overnight cultures were then diluted 1:100 into 50 ml YPAD or YPAD+1µg/ml FLC and grown at 30℃, 220 rpm for 5-6 hrs to OD_600_ of 0.5. Cells were harvested by centrifugation and frozen in liquid nitrogen. RNA were prepared according to the manufacturer’s instructions for the Qiagen RNeasy Mini kit (Qiagen, US) using the mechanical disruption method. Removal of DNA was performed with a DNase (Qiagen RNase-free DNase set, US) 1 hr incubation at room temperature on column. Three independent cultures of each strain were grown to provide three biological replicates for RNA-seq experiments.
ii) RNA-Seq: Library preparation was performed by SeqCenter (Pittsburgh, PA) using Illumina’s Stranded mRNA preparation and 10bp unique dual indices (UDI). Sequencing was done on a NovaSeq X Plus, producing 150bp paired end reads. Demultiplexing, quality control, and adapter trimming was performed with bcl-convert (v4.1.5) (<u>BCL Convert</u>).
iii) RNA-Seq data analysis: *C. albicans* transcriptome (SC5314_version_A21-s02-m09-r10_orf_coding, downloaded from http://www.candidagenome.org/download/sequence/C_albicans_SC5314/Assembly21/current/? C=S;O=A on 2023/08/17) was indexed using salmon (v1.10.2) [91]. All samples were quasi-mapped to transcriptome index using salmon resulting in quantification of reads mapped to each transcript. The output quantification files were imported into R (v4.1.2) using tximport (v1.22.0) [92] and DESeq2 (1.34.0) [93] was used to model gene expression. PCA analysis was performed using DESeq2 and used to identify any outliers amongst the replicates. We identified one wild-type control grown in FLC as an outlier and excluded this sample from all further analyses (Fig 5C). The DESeq2 ‘contrast’ wrapper was then used to estimate log_2_ fold changes for each mutant relative to the wild-type control in YPAD and YPAD+1µg/ml FLC conditions and identify differentially expressed genes. We also estimated log_2_ fold changes for each strain grown in FLC relative to the same strain in YPAD and identified differentially expressed genes. The threshold for differentially expressed genes was an absolute value log_2_ fold change ≥ 0.5 and adjusted p-value < 0.1 for heterozygous deletion mutants. Because the homozygous deletion mutant is predicted to have stronger effects on global gene expression, we used stricter thresholds for the homozygous deletion mutant of an absolute value log_2_ fold change ≥ 1 and adjusted p-value < 0.05. Differentially expressed genes in *ERG251* mutants or after FLC exposure are listed in supplementary tables.

##### Gene Ontology Analysis

GO slim mapper from *Candida* Genome Database (http://www.candidagenome.org/) [94] was conducted on the set of genes that were different expressed in *ERG251* mutants grown in YPAD relative to wild-type controls in YPAD. Process Ontology was performed for all three *ERG251* deletion mutants and output files are included in supplementary tables.

##### Rhodamine 6G efflux assay

Drug efflux was measured using an adapted protocol [29,47]. Strains were struck on YPAD agar from frozen stocks and incubated at 30℃ for 24 hrs. Recovered cells were inoculated into 2% dextrose YPAD or YPAD+1 µg/ml FLC. Cultures were incubated at 30℃, 220 rpm, for 16 hrs. Cultures were diluted 1:100 into fresh media of the same condition, then incubated 30℃, 220 rpm, for 3 hrs. Subcultures were harvested and washed once with room temperature PBS, then resuspended in PBS and incubated at 30℃ for 1 hr. Rhodamine 6G (Sigma) was added to a final concentration of 10 µg/ml. Cells were incubated at 30℃ for 1 hr. Following incubation, cells were washed twice with 4℃ PBS, then resuspended in room temperature PBS. Immediately, OD_600_ and baseline fluorescence were measured (excitation 344 nm, emission 555 nm) for 5 minutes in 1-minute intervals using a BioTek Synergy H1 plate reader. Following initial measurements, dextrose was added to a final concentration of 1%. Fluorescence was measured for 90 minutes in 2-minute intervals using a BioTek Synergy H1 plate reader. All strains were conducted in three independent replicates and tested with and without dextrose.

##### Growth curve assay

Strains were inoculated in 2% dextrose YPAD from frozen stocks and incubated at 30℃, 220 rpm for 16 hrs. Cultures were diluted in fresh 1% dextrose YPAD to a final OD_600_ of 0.01. Normalized cultures were diluted 1:10 into a 96-well NUNC plate containing 1% dextrose YPAD with or without drug. Cells were incubated at 30℃ in a BioTek Epoch 2 microplate spectrophotometer shaking in a double orbital (237rpm) with OD_600_ readings taken every 15 minutes for 48 hrs. Each isolate was conducted in triplicates.

##### Growth assay for FLC and farnesol

Strains were inoculated in 2% dextrose YPAD from frozen stocks and incubated at 30℃, 220 rpm for 16 hrs. Cultures were diluted in fresh 1% dextrose YPAD to a final OD_600_ of 0.01. Normalized cultures were diluted 1:10 into a 96-well NUNC plate containing 1% dextrose YPAD supplemented with or without drug. Both FLC and farnesol were diluted in a two-fold serial: FLC concentration (x-axis, 2x dilution for each column) ranged from 0 to 256µg/ml, and farnesol concentration (y-axis, 2x dilution for each row) ranged from 0 to 1000µM. Cells were incubated at 30℃ in a BioTek Epoch 2 microplate spectrophotometer shaking in a double orbital (237rpm) with OD_600_ readings taken every 15 minutes for 48hrs. Plates were conducted in triplicates. 48 hrs later, 10 µl cells from each well were plated onto YPAD agar plate to monitor viability, and plate images were taken after 24 hrs incubation at 30℃.

##### Relative fitness assay

Isolates were inoculated in 2% dextrose YPAD from frozen stocks and incubated at 30℃, 220 rpm, for 16hrs. Cultures were diluted in fresh 1% dextrose YPAD to a final OD_600_ of 0.01. Normalized cultures from the sample of interest and the fluorescent control strain (same fitness as the WT) were then combined at a 1:1 ratio, and the combined culture was diluted 1:10 into a 96-well NUNC plate containing 1% dextrose YPAD or YPAD+1 µg/ml FLC (initial OD_600_, N_0_=0.001). Cells were incubated at 30℃ in a BioTek Epoch 2 microplate spectrophotometer with double-orbital (237rpm) shaking. OD600 readings were taken every 15 minutes for 48 hrs to monitor cell growth and OD_600_ at the endpoint. 20 μl culture was removed from one of the triplicates for flow cytometry at 24 hrs. Cultures were diluted in PBS and 10,000 singlets were gated and analyzed at each time point using a Cytek Aurora flow cytometer (R0021). After 24 hrs the population reached the stationary phase and OD_600_ was about 1.3 (Nt), therefore there were a total 10 generations for the competition assay estimated using equation generations=[log10 (Nt/N0)]/0.3. Proportions of sample interest were indicated by the proportion of non-fluorescent cells while fluorescent control was indicated by blue fluorescent cells. All competitive assays were conducted in three independent replicates. Relative fitness was estimated using natural log regression analysis of the proportion of sample of interest and fluorescent control against the generations (10 generations): ln (proportion of sample of interest/proportion of fluorescent control)/generations.

##### Microdilution MIC and SMG assays

Isolates were inoculated in 2% dextrose YPAD from frozen stocks and incubated at 30 °C, 220 rpm, for 16 hrs. Cultures were diluted in fresh 1% dextrose YPAD to a final OD_600_ of 0.01. Normalized cultures were diluted 1:10 into 1% dextrose YPAD media containing either a two-fold serial dilution of drug or a no-drug control. Drug concentrations ranged from 0.5μg/ml to 256 μg/ml FLC and 0.0625 μg/ml to 32 μg/ml itraconazole and voriconazole. Triplicates of each isolate were set up using flat-bottom 96-well plates and incubated in a humidified chamber at 30°C. Cells were resuspended at the 24 hrs and 48 hrs time points and OD_600_ readings were taken using a BioTek Epoch 2 microplate spectrophotometer. The MIC_50_ of each strain was determined as the drug concentration at which ≥ 50% of growth was inhibited relative to the no-drug control at 24 hrs post-inoculation. The supra-MIC growth (SMG) was measured as the average growth above the MIC_50_ when standardized to the no-drug control at 48 hrs post-inoculation [9]. To measure the impact of Hsp90 inhibition, 2.5 µM radicicol (Cayman Chemicals) was added to the 1% dextrose YPAD in the microdilution MIC and SMG assay plate. To determine cell viability, 5µl was removed from the assay plate after the 48 hr time point and plated onto YPAD agar plates without any drugs. Plate images were taken after 24 hrs incubation at 30 °C.

##### Ploidy analysis (DNA-PI staining)

Cells were prepared as described previously [95]. Strains were inoculated in 2% dextrose YPAD from frozen stocks and incubated at 30℃, 220 rpm, for 16 hrs (cell density ∼1×10^7^ cells/ml). Cultures were spun down and the supernatant was removed. Cell pellets were resuspended in 70% ethanol, and then washed twice with 50 mM sodium citrate. Cells were then treated with RNAse A at 37℃ for at least 2 hrs, and then stained with 25 µg/ml propidium iodide (PI) at 37℃ in the dark for 16 hrs. Samples were diluted in 50 mM sodium citrate and at least 10,000 singlets were gated and analyzed using a Cytek Aurora flow cytometer (R0021). 488-nm lasers were used to excite the PI-staining and 618/24 filters were used to detect the PI-staining emission signals. Data were analyzed using FlowJo v10.8.1.

##### Illumina whole genome sequencing

Genomic DNA was isolated using a phenol-chloroform extraction as described previously [96]. Libraries were prepared using the Illumina DNA Prep kit and IDT 10bp UDI indices, and sequenced on an Illumina NextSeq 2000, producing 2×151bp reads. Demultiplexing, quality control, and adapter trimming were performed with bcl-convert (https://support.illumina.com/sequencing/sequencing_software/bcl-convert.html)(v3.9.3).

Adapter and quality trimming were performed with BBDuk (BBTools v38.94) [97]. Trimmed reads were aligned to the *C. albicans* reference genome (SC5314_version_A21-s02-m09-r08) using BWA-MEM (v0.7.17) with default parameters [98,99]. Aligned reads were sorted, duplicate reads were marked and the resulting BAM file was indexed with Samtools (v1.10) [99]. Quality of trimmed FASTQ and BAM files was assessed for all strains with FastQC (v0.11.7), Qualimap (v2.2.2-dev) and MultiQC (v1.16) [100–102].

##### Visualization of whole genome sequencing data

Chromosomal copy number changes were visualized using the Yeast Mapping Analysis Pipeline (YMAP v1.0). Aligned BAM files were uploaded to YMAP and read depth was determined and plotted as a function of chromosome position using the reference genome *C. albicans* SC5314 (A21-s02-m09-r08). Read depth was corrected for GC-content and chromosome-end bias. WGS data were plotted as the log2 ratio and converted to chromosome copy number (y-axis, 1-4 copies) as a function of chromosome position (x-axis, Chr1-ChrR) using the Yeast Mapping Pipeline (YMAP) [103]. The baseline chromosome copy number (ploidy) was determined by flow cytometry (S1 Table). Haplotypes are indicated by color: gray is heterozygous (AB), magenta is homozygous B, and cyan is homozygous A.

##### Variant calling

De novo variant calling and preliminary filtering were performed with Mutect2 and FilterMutectCalls (GATK v4.1.2), both with default parameters as previously described [104]. Variant calling was run separately for 3 groups of strains corresponding to different progenitors. The first group called the Sn152 progenitor as “normal” and Sn152-evolved strains as “tumor”. The second group called BWP17 as “normal” and BWP17-evolved strains as “tumor”. The third group called P75063 as “normal” and P75063-evolved strains as “tumor”. Additional VCF filtering was performed with bcftools (v1.17) [99]. Individual VCF files were subset to remove the progenitor strain, and filtered for calls with a quality status of “PASS”. A merged VCF file was created for each progenitor group. Merged VCF files were subset to exclude repeat regions (as marked in the SC5314 A21-s02-m09-r08 GFF) and 5000 bp subtelomeric regions, and additional hard filtering was performed (minimum 5 supporting reads, at least one supporting read in both forward and reverse direction, minimum alternate allele frequency of 0.2 for diploid, single colony cultures). Identical variants found in at least half of all progeny were considered to be present in the progenitor strain and were removed [10]. Variants were annotated with SnpEff (v5.0e, database built from SC5314 version A21-s02-m09-r08, with alternate yeast nuclear codon table) and visually verified in IGV [105,106]. All variants of *ERG251* were compiled into S13 Table.

### Murine model

*C. albicans* strains were serially passaged three times in YPD broth, grown in a shaking incubator at 30°C for 16-24h at each passage. To prepare *C. albicans* for infection, yeast cells were collected by centrifugation, washed in PBS, and counted using a hemocytometer. Male, 5-6 weeks old ICR mice (Envigo) were infected with 2×10^5^ *C. albicans* yeast cells via the lateral tail vein. Mice were monitored three times daily for survival for 21 days. Moribund mice were humanely euthanized.

### Ethics statement

The mouse experiments were approved by the Institutional Animal Care and Use Committee of the Lundquist Institute for Biomedical Innovation at Harbor-UCLA Medical Center.

### Data availability

All whole genome sequences and RNA sequences are available in the Sequence Read Archive repository under BioProject PRJNA1068093 and BioProject PRJNA1068582.

## ACKNOWLEDGEMENTS

We are grateful to Berman lab, Cowen lab and Köhler lab for the plasmids used for strain engineering: pMG2120, pLC605 and pJK863. We thank Luke Dragseth, Maicy Vossen and Hanaa Alhosawi for technical assistance with the evolution and sequencing of some of the evolved strains where *ERG251* mutations were initially identified. We are grateful to Petra Vande Zande for helpful discussions and feedback on the manuscript. Funding for this work was provided by the National Institutes of Health (R01 AI143689) and Burroughs Wellcome Fund Investigator in the Pathogenesis of Infectious Diseases Award (#1020388) to AS, the Swanson-Holcomb Undergraduate Research Fund at Gustavus Adolphus College to TB and LSB, and the First Year Research Experience Award at Gustavus Adolphus College to BH.

## AUTHOR CONTRIBUTIONS

**Conceptualization:** Xin Zhou, Laura S. Burrack, Anna Selmecki

**Investigation:** Xin Zhou, Audrey Hilk, Norma V. Solis, Bode M. Hogan, Tessa Bierbaum

**Formal analysis**: Xin Zhou, Audrey Hilk

**Methodology:** Xin Zhou, Laura Burrack, Norma V. Solis, Anna Selmecki

**Project administration**: Xin Zhou, Anna Selmecki

**Resources**: Scott G. Filler, Laura S. Burrack, Anna Selmecki

**Supervision**: Scott G. Filler, Laura S. Burrack, Anna Selmecki

**Writing-original draft:** Xin Zhou

**Writing-Review & Editing**: Xin Zhou, Laura Burrack, Anna Selmecki

## SUPPLEMENTARY INFORMATION

**Fig S1.**
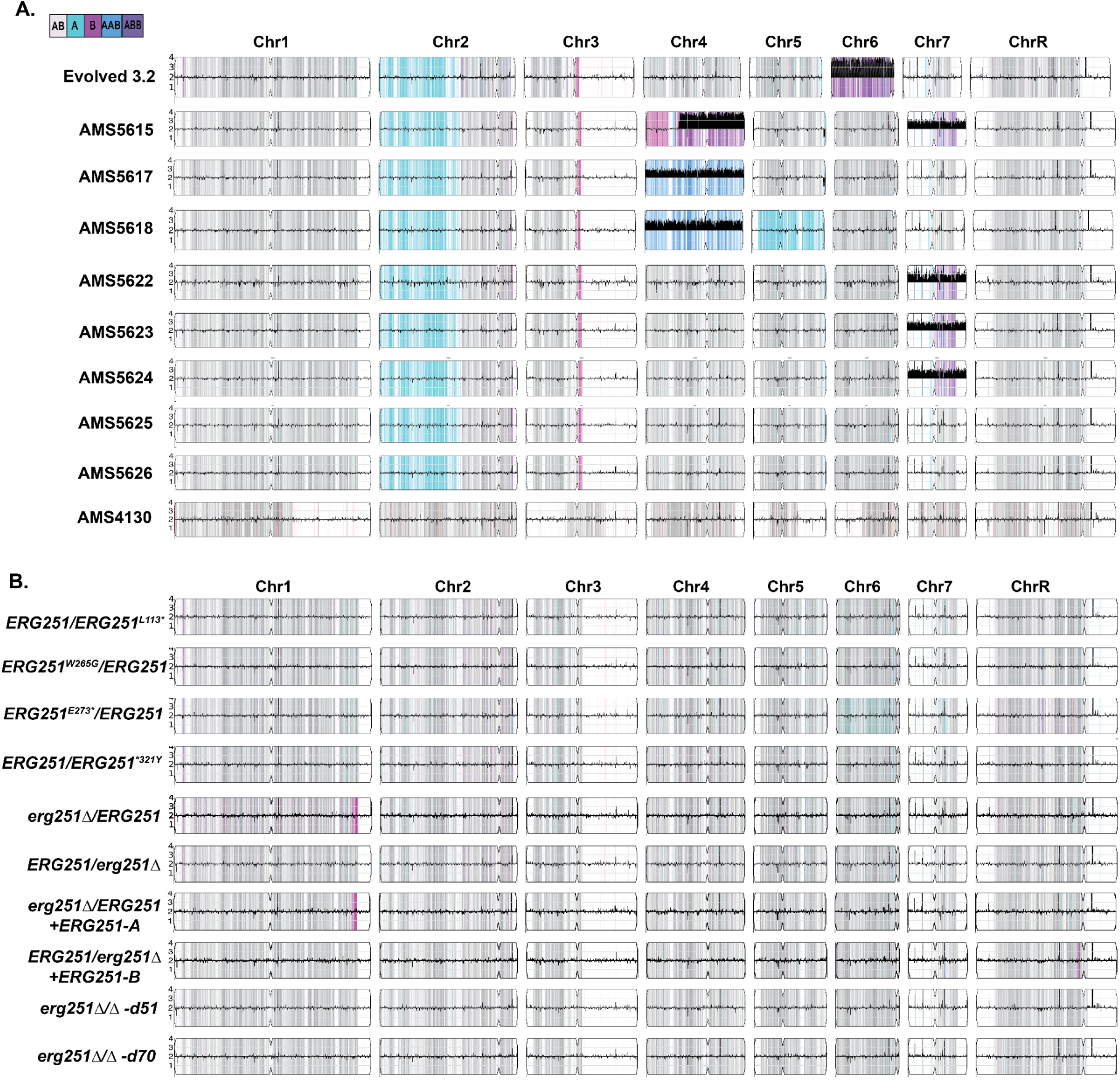
Whole genome sequencing analysis of FLC-evolved and engineered strains. **A.** *De novo* point mutations in *ERG251* often occur together with other aneuploidies. Representative whole genome sequencing (WGS) data of the FLC-evolved strains from Table 1: Evolved 3.2, AMS5615, AMS5617, AMS5618, AMS5622, AMS5623, AMS5624, AMS5625, AMS5626 and AMS4130 which acquired point mutations on *ERG251* during FLC evolution. **B.** The engineered *ERG251* mutants remain euploid. WGS data for all *ERG251* mutations engineered into the euploid SC5314 genetic background: the *ERG251* heterozygous point mutants (L113*, W265G, E273*, and *321Y), both heterozygous deletion strains of *ERG251*, two strains with complementation of the heterozygous deletion, and two independent homozygous deletions of *ERG251* (d51 and d70). **A&B** WGS data are plotted as the log2 ratio and converted to chromosome copy number (y-axis, 1-4 copies) as a function of chromosome position (x-axis, Chr1-ChrR). Haplotypes are indicated by color: gray is heterozygous (AB), magenta is homozygous B, and cyan is homozygous A. The baseline ploidy was determined by propidium iodide staining (S1 Table).

**Fig S2.**
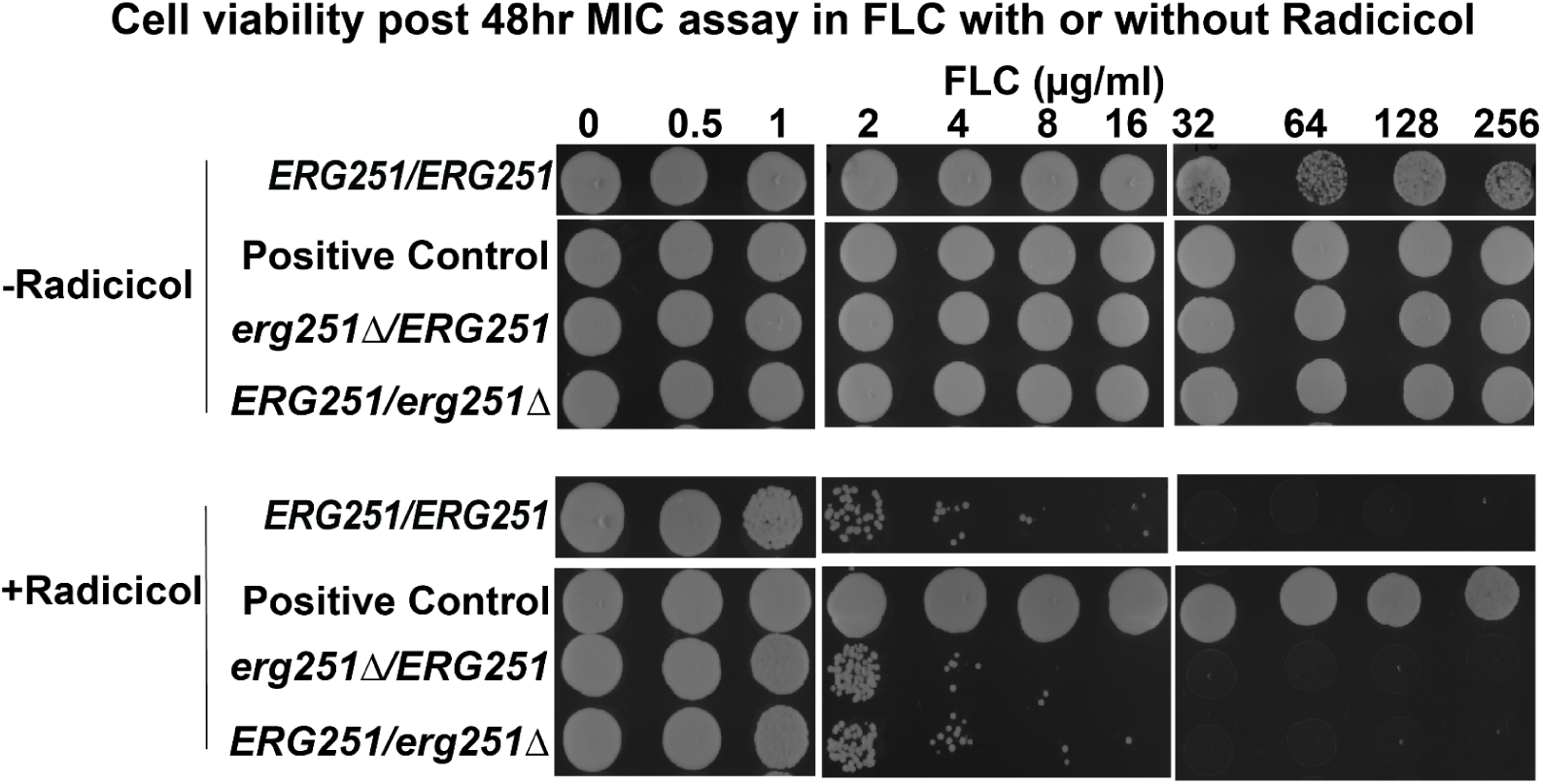
**Radicicol, an Hsp90 inhibitor, blocks Erg251-driven tolerance and makes fluconazole fungicidal.** Cells from the MIC assay at 48 hr in Fig 1D, with or without radicicol, were plated for viability on YPAD agar plates and imaged after 24 hr incubation. Wildtype SC5314 (*ERG251/ERG251*), a positive control strain known to be resistant to fluconazole (FLC), and both heterozygous deletion mutants of *ERG251* were tested. At least three biological replicates were performed.

**Fig S3.**
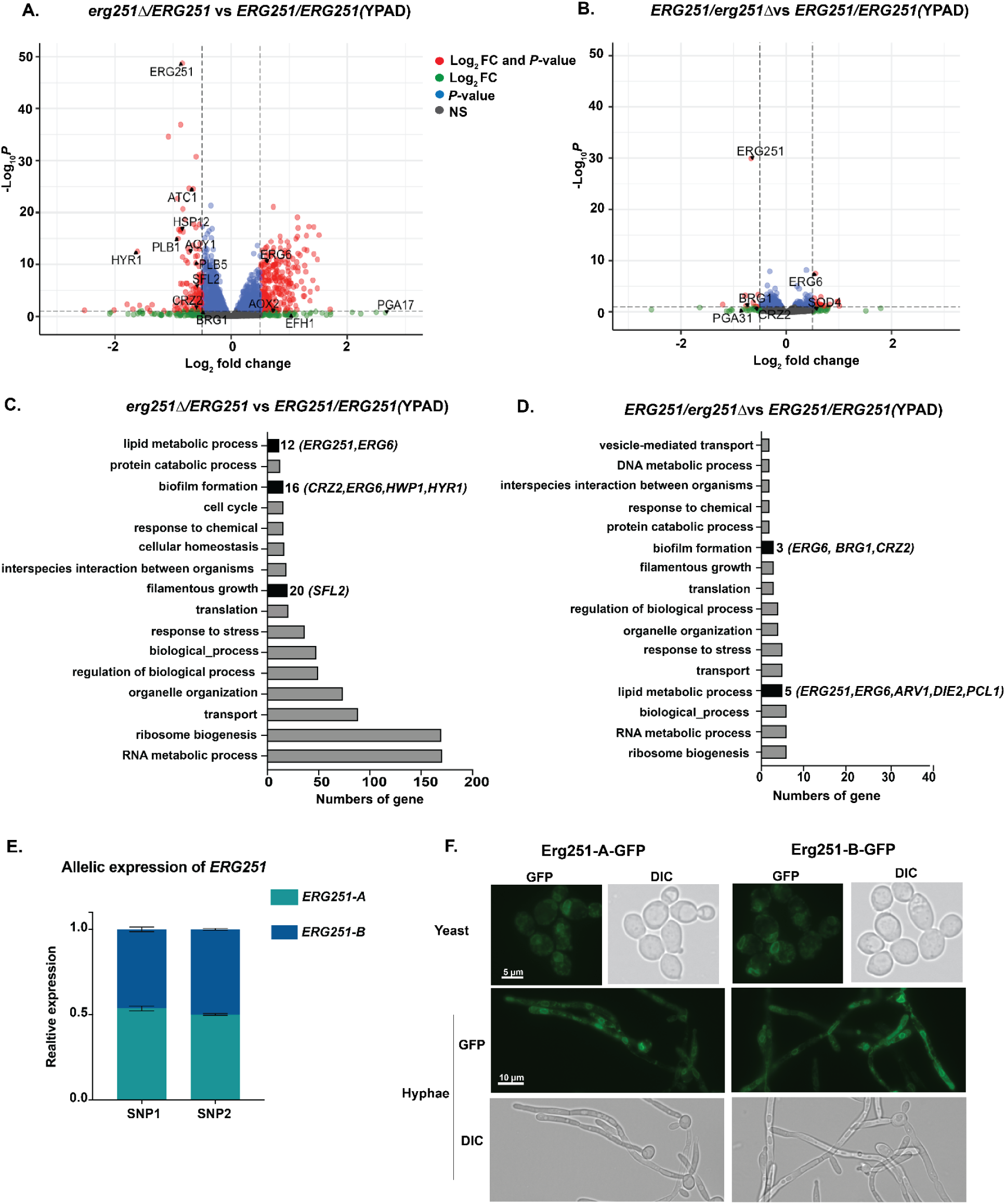
Heterozygous deletion of *ERG251-A* leads to a transcriptional response in filamentation regulation. Volcano plots for differentially expressed genes (log2 fold change ≥ 0.5 or ≤ -0.5 and adjusted *p*-value < 0.1) in the heterozygous mutants **(A)** *erg251Δ/ERG251* and **(B)** *ERG251/erg251Δ* in YPAD compared to the wildtype *ERG251/ERG251* in YPAD. Both the fold change and *p*-value are indicated. **C&D.** Gene Ontology (GO) terms for genes differentially expressed in (C, S7 Table) *erg251Δ/ERG251* in YPAD and (D, S8 Table) *ERG251/erg251Δ* in YPAD compared to *ERG251/ERG251* in YPAD. **E.** Relative expression of *ERG251-A* and *ERG251-B* in the SC5314 background in YPAD. Relative expression was estimated using allelic RNA reads compared to overall reads at the two loci with polymorphisms in the *ERG251* gene (indicated as SNP1 and SNP2 above). Values are mean ± SEM calculated from three biological replicates. **F.** Subcellular localization of Erg251-A-GFP and Erg251-B-GFP in yeast and hyphal inducing conditions in SC5314 background. Yeast: scale bar, 5 μm; hyphae: scale bar, 10 μm.

**Fig S4.**
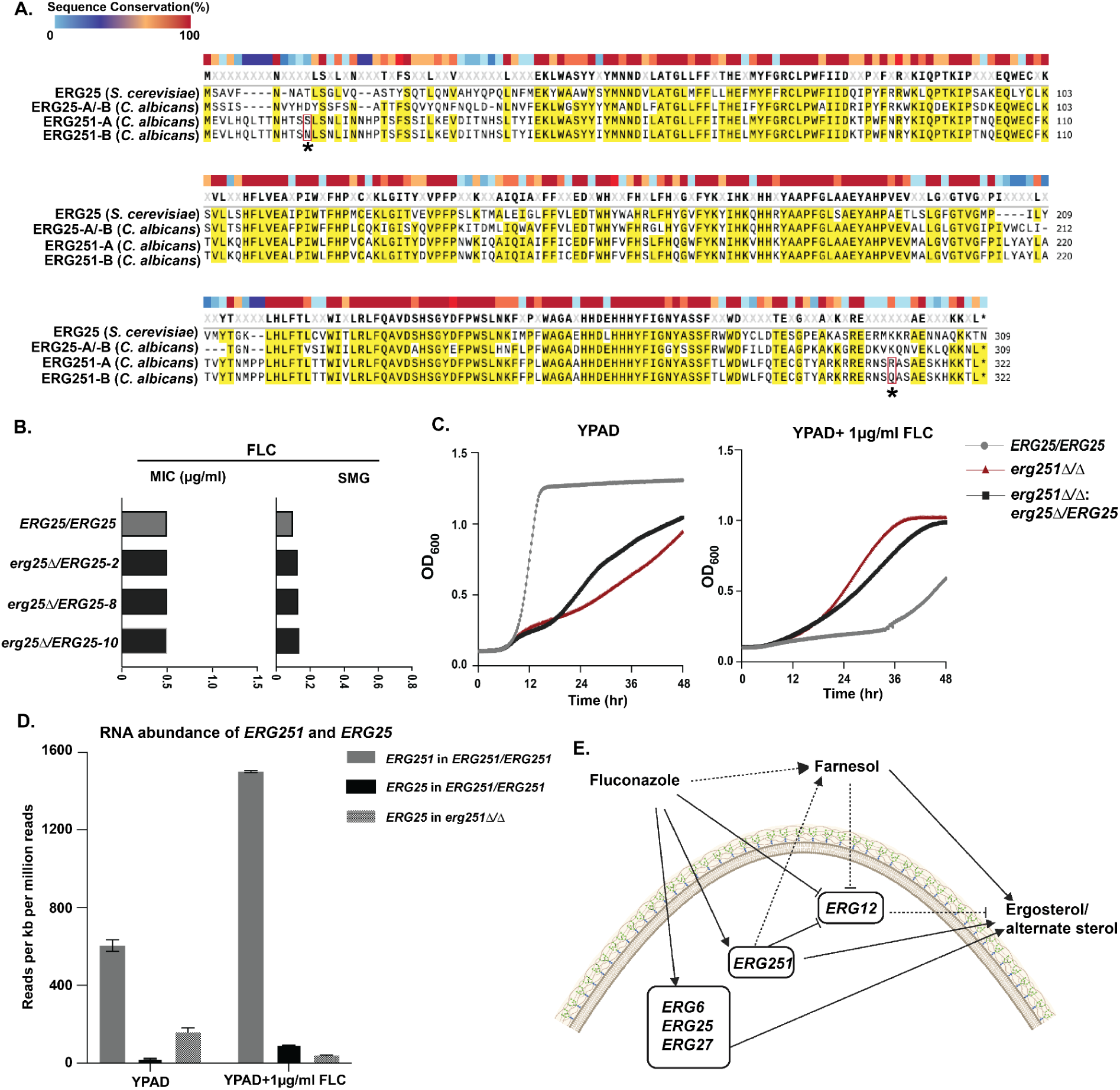
Erg251 is the major methyl sterol oxidase controlling drug susceptibility compared to its paralog Erg25. (**A**) Multiple sequence alignment for *ERG251-A*, *ERG251-B*, and *ERG25*-A/-B (no SNPs between A and B) from *C. albicans* and *ERG25* from *S. cerevisiae,* with yellow highlighting similarity among all four proteins. Colored blocks on the top indicate the sequence conservation. Asterisks (*) and red boxes indicate the locus of non-synonymous variation between *ERG251-A* and *ERG251-B* in *C. albicans.* **B.** FLC susceptibility determined by liquid microbroth dilution at 24hr MIC (left, µg/ml) and 48hr SMG (right, tolerance) in FLC for three *ERG25* heterozygous deletion mutants (*ERG25/erg25Δ-2, -8* and *-10*) in the SC5314 background with SC5314 (*ERG25/ERG25*) as the control. **C.** 48hr growth curve analysis of *erg25* heterozygous deletion strain in *erg251Δ/Δ* background (*erg251Δ/Δ: ERG25/erg25Δ*) in YPAD (left) and YPAD+1µg/ml FLC (right) with SC5314 (*ERG25/ERG25*) and *erg251Δ/Δ* as the controls. The initial cell densities were OD_600_ of 0.001. MIC and SMG are not measurable for *erg251Δ/Δ* or *erg251Δ/Δ: ERG25/erg25Δ* given growth defects in YPAD. **B&C**: Minimum of three biological replicates were performed. **D.** RNA abundance of *ERG251* and *ERG25* in SC5314 (*ERG251/ERG251*), and *ERG25* in *erg251Δ/Δ*. RNA reads were normalised to transcript length and total RNA reads. Values are mean ± SEM calculated from three biology replicates. **E.** Predicted model for how FLC and farnesol impact the expression of *ERG* genes. In the wildtype, low concentrations of FLC promote the expression of most *ERG* genes, including *ERG6*, *ERG251*, *ERG25*, *ERG11* and *ERG27*, leading to the upregulation of ergosterol or/and alternate sterol biosynthesis. However, both low concentrations of FLC and Erg251 pose a negative regulation on Erg12, which may be achieved via farnesol which we predict inhibits *ERG12* [107]. Dashed lines indicate predicted relationships. Figure created in BioRender.com.

## SUPPLEMENTARY TABLES

**S1 Table**. Strains used in this study.

**S2 Table**. Differentially expressed genes in *erg251Δ/Δ* in YPAD compared to wildtype in YPAD.

**S3 Table.** GO term analysis for differentially expressed genes in *erg251Δ/Δ* in YPAD compared to wildtype in YPAD.

**S4 Table.** Differentially expressed GPI genes in *erg251Δ/Δ* in YPAD compared to wildtype in YPAD.

**S5 Table.** Differentially expressed genes in *erg251Δ/ERG251* in YPAD compared to wildtype in YPAD.

**S6 Table**. Differentially expressed genes in *ERG251/erg251Δ* in YPAD compared to wildtype in YPAD.

**S7 Table**. GO term for differentially expressed genes in *erg251Δ/ERG251* in YPAD compared to wildtype in YPAD.

**S8 Table**. GO term for differentially expressed genes in *ERG251/erg251Δ* in YPAD compared to wildtype in YPAD.

**S9 Table**. Differentially expressed genes in *erg251Δ/ERG251* in FLC compared to wildtype in FLC.

**S10 Table**. Differentially expressed genes in *ERG251/erg251Δ* in FLC compared to wildtype in FLC.

**S11 Table**. Differentially expressed genes in *erg251Δ/Δ* in FLC compared to wildtype in FLC.

**S12 Table**. Primers used in this study.

**S13 Table**. *ERG251* SNPs from all FLC-evovled strains.

## REFERENCES

1. Pfaller MA, Diekema DJ, Turnidge JD, Castanheira M, Jones RN. Twenty Years of the SENTRY Antifungal Surveillance Program: Results for Candida Species From 1997–2016. Open Forum Infect Dis. 2019;6: S79–S94.

2. Pfaller MA. Antifungal drug resistance: mechanisms, epidemiology, and consequences for treatment. Am J Med. 2012;125: S3–13.

3. Perea S, López-Ribot JL, Kirkpatrick WR, McAtee RK, Santillán RA, Martínez M, et al. Prevalence of molecular mechanisms of resistance to azole antifungal agents in Candida albicans strains displaying high-level fluconazole resistance isolated from human immunodeficiency virus-infected patients. Antimicrob Agents Chemother. 2001;45: 2676–2684.

4. Pappas PG, Kauffman CA, Andes DR, Clancy CJ, Marr KA, Ostrosky-Zeichner L, et al. Clinical Practice Guideline for the Management of Candidiasis: 2016 Update by the Infectious Diseases Society of America. Clin Infect Dis. 2015;62: e1–e50.

5. Andes DR, Safdar N, Baddley JW, Playford G, Reboli AC, Rex JH, et al. Impact of treatment strategy on outcomes in patients with candidemia and other forms of invasive candidiasis: a patient-level quantitative review of randomized trials. Clin Infect Dis. 2012;54: 1110–1122.

6. Cowen LE. The evolution of fungal drug resistance: modulating the trajectory from genotype to phenotype. Nat Rev Microbiol. 2008;6: 187–198.

7. Berman J, Krysan DJ. Drug resistance and tolerance in fungi. Nat Rev Microbiol. 2020;18: 319–331.

8. Sanglard D. Emerging Threats in Antifungal-Resistant Fungal Pathogens. Front Med. 2016;3: 11.

9. Rosenberg A, Ene IV, Bibi M, Zakin S, Segal ES, Ziv N, et al. Antifungal tolerance is a subpopulation effect distinct from resistance and is associated with persistent candidemia. Nat Commun. 2018;9: 2470.

10. Todd RT, Soisangwan N, Peters S, Kemp B, Crooks T, Gerstein A, et al. Antifungal Drug Concentration Impacts the Spectrum of Adaptive Mutations in Candida albicans. Mol Biol Evol. 2023;40: msad009.

11. White TC, Marr KA, Bowden RA. Clinical, cellular, and molecular factors that contribute to antifungal drug resistance. Clin Microbiol Rev. 1998;11: 382–402.

12. Shapiro RS, Robbins N, Cowen LE. Regulatory circuitry governing fungal development, drug resistance, and disease. Microbiol Mol Biol Rev. 2011;75: 213–267.

13. Cowen LE, Steinbach WJ. Stress, drugs, and evolution: the role of cellular signaling in fungal drug resistance. Eukaryot Cell. 2008;7: 747–764.

14. Kelly SL, Lamb DC, Kelly DE, Manning NJ, Loeffler J, Hebart H, et al. Resistance to fluconazole and cross-resistance to amphotericin B in Candida albicans from AIDS patients caused by defective sterol delta5,6-desaturation. FEBS Lett. 1997;400: 80–82.

15. Lupetti A, Danesi R, Campa M, Del Tacca M, Kelly S. Molecular basis of resistance to azole antifungals. Trends Mol Med. 2002;8: 76–81.

16. Sanglard D, Kuchler K, Ischer F, Pagani JL, Monod M, Bille J. Mechanisms of resistance to azole antifungal agents in Candida albicans isolates from AIDS patients involve specific multidrug transporters. Antimicrob Agents Chemother. 1995;39: 2378–2386.

17. Revie NM, Iyer KR, Robbins N, Cowen LE. Antifungal drug resistance: evolution, mechanisms and impact. Curr Opin Microbiol. 2018;45: 70–76.

18. Shrivastava Manjari, Kouyoumdjian Gaëlle S., Kirbizakis Eftyhios, Ruiz Daniel, Henry Manon, Vincent Antony T., et al. The Adr1 transcription factor directs regulation of the ergosterol pathway and azole resistance in Candida albicans. MBio. 2023;14: e01807–23.

19. Blosser SJ, Merriman B, Grahl N, Chung D, Cramer RA. Two C4-sterol methyl oxidases (Erg25) catalyse ergosterol intermediate demethylation and impact environmental stress adaptation in Aspergillus fumigatus. Microbiology. 2014;160: 2492–2506.

20. Lu H, Li W, Whiteway M, Wang H, Zhu S, Ji Z, et al. A Small Molecule Inhibitor of Erg251 Makes Fluconazole Fungicidal by Inhibiting the Synthesis of the 14α-Methylsterols. MBio. 2023;14: e0263922.

21. Bhattacharya Somanon, Esquivel Brooke D., White Theodore C. Overexpression or Deletion of Ergosterol Biosynthesis Genes Alters Doubling Time, Response to Stress Agents, and Drug Susceptibility in Saccharomyces cerevisiae. MBio. 2018;9: 10.1128/mbio.01291–18.

22. Martel CM, Parker JE, Bader O, Weig M, Gross U, Warrilow AGS, et al. Identification and characterization of four azole-resistant erg3 mutants of Candida albicans. Antimicrob Agents Chemother. 2010;54: 4527–4533.

23. Jordá T, Puig S. Regulation of Ergosterol Biosynthesis in Saccharomyces cerevisiae. Genes . 2020;11. doi:10.3390/genes11070795

24. Hornby JM, Jensen EC, Lisec AD, Tasto JJ, Jahnke B, Shoemaker R, et al. Quorum sensing in the dimorphic fungus Candida albicans is mediated by farnesol. Appl Environ Microbiol. 2001;67: 2982–2992.

25. Hornby JM. Quorum sensing and the regulation of morphology in the dimorphic fungus Candida albicans. 2003. Available: https://search.proquest.com/openview/0b2bfd0efe29f67bc0222719435cd8b2/1?pq-origsite=gscholar&cbl=18750&diss=y

26. Ramage G, Saville SP, Wickes BL, López-Ribot JL. Inhibition of Candida albicans biofilm formation by farnesol, a quorum-sensing molecule. Appl Environ Microbiol. 2002;68: 5459–5463.

27. Sanglard D, Ischer F, Parkinson T, Falconer D, Bille J. Candida albicans mutations in the ergosterol biosynthetic pathway and resistance to several antifungal agents. Antimicrob Agents Chemother. 2003;47: 2404–2412.

28. Jensen-Pergakes KL, Kennedy MA, Lees ND, Barbuch R, Koegel C, Bard M. Sequencing, disruption, and characterization of the Candida albicans sterol methyltransferase (ERG6) gene: drug susceptibility studies in erg6 mutants. Antimicrob Agents Chemother. 1998;42: 1160–1167.

29. Vale-Silva LA, Coste AT, Ischer F, Parker JE, Kelly SL, Pinto E, et al. Azole resistance by loss of function of the sterol Δ^5^,^6^-desaturase gene (ERG3) in Candida albicans does not necessarily decrease virulence. Antimicrob Agents Chemother. 2012;56: 1960–1968.

30. Young LY, Hull CM, Heitman J. Disruption of ergosterol biosynthesis confers resistance to amphotericin B in Candida lusitaniae. Antimicrob Agents Chemother. 2003;47: 2717–2724.

31. Pinjon E, Moran GP, Jackson CJ, Kelly SL, Sanglard D, Coleman DC, et al. Molecular mechanisms of itraconazole resistance in Candida dubliniensis. Antimicrob Agents Chemother. 2003;47: 2424–2437.

32. Morio F, Pagniez F, Lacroix C, Miegeville M, Le Pape P. Amino acid substitutions in the Candida albicans sterol Δ5,6-desaturase (Erg3p) confer azole resistance: characterization of two novel mutants with impaired virulence. J Antimicrob Chemother. 2012;67: 2131–2138.

33. Ksiezopolska E, Schikora-Tamarit MÀ, Beyer R, Nunez-Rodriguez JC, Schüller C, Gabaldón T. Narrow mutational signatures drive acquisition of multidrug resistance in the fungal pathogen Candida glabrata. Curr Biol. 2021;31: 5314–5326.e10.

34. Vandeputte P, Tronchin G, Larcher G, Ernoult E, Bergès T, Chabasse D, et al. A nonsense mutation in the ERG6 gene leads to reduced susceptibility to polyenes in a clinical isolate of Candida glabrata. Antimicrob Agents Chemother. 2008;52: 3701–3709.

35. Carolus H, Sofras D, Boccarella G, Sephton-Clark P, Romero CL, Vergauwen R, et al. Acquired amphotericin B resistance and fitness trade-off compensation in Candida auris. Research Square. 2023. doi:10.21203/rs.3.rs-3621420/v1

36. Gao J, Wang H, Li Z, Wong AH-H, Wang Y-Z, Guo Y, et al. Candida albicans gains azole resistance by altering sphingolipid composition. Nat Commun. 2018;9: 1–14.

37. Rybak JM, Dickens CM, Parker JE, Caudle KE, Manigaba K, Whaley SG, et al. Loss of C-5 Sterol Desaturase Activity Results in Increased Resistance to Azole and Echinocandin Antifungals in a Clinical Isolate of Candida parapsilosis. Antimicrob Agents Chemother. 2017;61. doi:10.1128/AAC.00651-17

38. Kennedy MA, Johnson TA, Lees ND, Barbuch R, Eckstein JA, Bard M. Cloning and sequencing of the Candida albicans C-4 sterol methyl oxidase gene (ERG25) and expression of an ERG25 conditional lethal mutation in Saccharomyces cerevisiae. Lipids. 2000;35: 257–262.

39. Kim SH, Steere L, Zhang Y-K, McGregor C, Hahne C, Zhou Y, et al. Inhibiting C-4 Methyl Sterol Oxidase with Novel Diazaborines to Target Fungal Plant Pathogens. ACS Chem Biol. 2022;17: 1343–1350.

40. Kodedová M, Sychrová H. Changes in the Sterol Composition of the Plasma Membrane Affect Membrane Potential, Salt Tolerance and the Activity of Multidrug Resistance Pumps in Saccharomyces cerevisiae. PLoS One. 2015;10: e0139306.

41. Li Y, Dai M, Zhang Y, Lu L. The sterol C-14 reductase Erg24 is responsible for ergosterol biosynthesis and ion homeostasis in Aspergillus fumigatus. Appl Microbiol Biotechnol. 2021;105: 1253–1268.

42. Gupta SS, Ton V-K, Beaudry V, Rulli S, Cunningham K, Rao R. Antifungal activity of amiodarone is mediated by disruption of calcium homeostasis. J Biol Chem. 2003;278: 28831–28839.

43. Barreto L, Canadell D, Petrezsélyová S, Navarrete C, Maresová L, Peréz-Valle J, et al. A genomewide screen for tolerance to cationic drugs reveals genes important for potassium homeostasis in Saccharomyces cerevisiae. Eukaryot Cell. 2011;10: 1241–1250.

44. Luna-Tapia A, Peters BM, Eberle KE, Kerns ME, Foster TP, Marrero L, et al. ERG2 and ERG24 Are Required for Normal Vacuolar Physiology as Well as Candida albicans Pathogenicity in a Murine Model of Disseminated but Not Vaginal Candidiasis. Eukaryot Cell. 2015;14: 1006–1016.

45. Kim SH, Iyer KR, Pardeshi L, Muñoz JF, Robbins N, Cuomo CA, et al. Genetic Analysis of Candida auris Implicates Hsp90 in Morphogenesis and Azole Tolerance and Cdr1 in Azole Resistance. MBio. 2019;10. doi:10.1128/mBio.02529-18

46. Robbins N, Cowen LE. Roles of Hsp90 in Candida albicans morphogenesis and virulence. Curr Opin Microbiol. 2023;75: 102351.

47. Maesaki S, Marichal P, Vanden Bossche H, Sanglard D, Kohno S. Rhodamine 6G efflux for the detection of CDR1-overexpressing azole-resistant Candida albicans strains. J Antimicrob Chemother. 1999;44: 27–31.

48. Mehmood A, Liu G, Wang X, Meng G, Wang C, Liu Y. Fungal Quorum-Sensing Molecules and Inhibitors with Potential Antifungal Activity: A Review. Molecules. 2019;24. doi:10.3390/molecules24101950

49. Nickerson KW, Atkin AL, Hornby JM. Quorum sensing in dimorphic fungi: farnesol and beyond. Appl Environ Microbiol. 2006;72: 3805–3813.

50. Yu L-H, Wei X, Ma M, Chen X-J, Xu S-B. Possible inhibitory molecular mechanism of farnesol on the development of fluconazole resistance in Candida albicans biofilm. Antimicrob Agents Chemother. 2012;56: 770–775.

51. Song J, Liu X, Li R. Sphingolipids: Regulators of azole drug resistance and fungal pathogenicity. Mol Microbiol. 2020;114: 891–905.

52. Kadosh D, Johnson AD. Induction of the Candida albicans filamentous growth program by relief of transcriptional repression: a genome-wide analysis. Mol Biol Cell. 2005;16: 2903–2912.

53. Villa S, Hamideh M, Weinstock A, Qasim MN, Hazbun TR, Sellam A, et al. Transcriptional control of hyphal morphogenesis in Candida albicans. FEMS Yeast Res. 2020;20. doi:10.1093/femsyr/foaa005

54. Znaidi S, Nesseir A, Chauvel M, Rossignol T, d’Enfert C. A comprehensive functional portrait of two heat shock factor-type transcriptional regulators involved in Candida albicans morphogenesis and virulence. PLoS Pathog. 2013;9: e1003519.

55. Carbrey JM, Cormack BP, Agre P. Aquaporin in Candida: characterization of a functional water channel protein. Yeast. 2001;18: 1391–1396.

56. Gong Y, Li T, Yu C, Sun S. Candida albicans Heat Shock Proteins and Hsps-Associated Signaling Pathways as Potential Antifungal Targets. Front Cell Infect Microbiol. 2017;7: 520.

57. Yan L, Li M, Cao Y, Gao P, Cao Y, Wang Y, et al. The alternative oxidase of Candida albicans causes reduced fluconazole susceptibility. J Antimicrob Chemother. 2009;64: 764–773.

58. Martchenko M, Alarco A-M, Harcus D, Whiteway M. Superoxide dismutases in Candida albicans: transcriptional regulation and functional characterization of the hyphal-induced SOD5 gene. Mol Biol Cell. 2004;15: 456–467.

59. Burgain A, Tebbji F, Khemiri I, Sellam A. Metabolic Reprogramming in the Opportunistic Yeast Candida albicans in Response to Hypoxia. mSphere. 2020;5. doi:10.1128/mSphere.00913-19

60. Plaine A, Walker L, Da Costa G, Mora-Montes HM, McKinnon A, Gow NAR, et al. Functional analysis of Candida albicans GPI-anchored proteins: roles in cell wall integrity and caspofungin sensitivity. Fungal Genet Biol. 2008;45: 1404–1414.

61. Richard M, de Groot P, Courtin O, Poulain D, Klis F, Gaillardin C. GPI7 affects cell-wall protein anchorage in Saccharomyces cerevisiae and Candida albicans. Microbiology. 2002;148: 2125–2133.

62. Victoria GS, Yadav B, Hauhnar L, Jain P, Bhatnagar S, Komath SS. Mutual co-regulation between GPI-N-acetylglucosaminyltransferase and ergosterol biosynthesis in Candida albicans. Biochem J. 2012;443: 619–625.

63. Yadav B, Bhatnagar S, Ahmad MF, Jain P, Pratyusha VA, Kumar P, et al. First step of glycosylphosphatidylinositol (GPI) biosynthesis cross-talks with ergosterol biosynthesis and Ras signaling in Candida albicans. J Biol Chem. 2014;289: 3365–3382.

64. Doedt T, Krishnamurthy S, Bockmühl DP, Tebarth B, Stempel C, Russell CL, et al. APSES proteins regulate morphogenesis and metabolism in Candida albicans. Mol Biol Cell. 2004;15: 3167–3180.

65. White SJ, Rosenbach A, Lephart P, Nguyen D, Benjamin A, Tzipori S, et al. Self-regulation of Candida albicans population size during GI colonization. PLoS Pathog. 2007;3: e184.

66. MacPherson S, Akache B, Weber S, De Deken X, Raymond M, Turcotte B. Candida albicans zinc cluster protein Upc2p confers resistance to antifungal drugs and is an activator of ergosterol biosynthetic genes. Antimicrob Agents Chemother. 2005;49: 1745–1752.

67. Silver PM, Oliver BG, White TC. Role of Candida albicans transcription factor Upc2p in drug resistance and sterol metabolism. Eukaryot Cell. 2004;3: 1391–1397.

68. Lv Q-Z, Yan L, Jiang Y-Y. The synthesis, regulation, and functions of sterols in Candida albicans: Well-known but still lots to learn. Virulence. 2016;7: 649–659.

69. Revie NM, Iyer KR, Maxson ME, Zhang J, Yan S, Fernandes CM, et al. Targeting fungal membrane homeostasis with imidazopyrazoindoles impairs azole resistance and biofilm formation. Nat Commun. 2022;13: 1–20.

70. Pierson CA, Eckstein J, Barbuch R, Bard M. Ergosterol gene expression in wild-type and ergosterol-deficient mutants of Candidaalbicans. Med Mycol. 2004;42: 385–389.

71. Veen M, Stahl U, Lang C. Combined overexpression of genes of the ergosterol biosynthetic pathway leads to accumulation of sterols in Saccharomyces cerevisiae. FEMS Yeast Res. 2003;4: 87–95.

72. Crawford AC, Lehtovirta-Morley LE, Alamir O, Niemiec MJ, Alawfi B, Alsarraf M, et al. Biphasic zinc compartmentalisation in a human fungal pathogen. PLoS Pathog. 2018;14: e1007013.

73. Sanchez AA, Johnston DA, Myers C, Edwards JE Jr, Mitchell AP, Filler SG. Relationship between Candida albicans virulence during experimental hematogenously disseminated infection and endothelial cell damage in vitro. Infect Immun. 2004;72: 598–601.

74. Kukurudz RJ, Chapel M, Wonitowy Q, Adamu Bukari A-R, Sidney B, Sierhuis R, et al. Acquisition of cross-azole tolerance and aneuploidy in Candida albicans strains evolved to posaconazole. G3 . 2022;12. doi:10.1093/g3journal/jkac156

75. Yang Feng, Scopel Eduardo F. C., Li Hao, Sun Liu-liu, Kawar Nora, Cao Yong-bing, et al. Antifungal Tolerance and Resistance Emerge at Distinct Drug Concentrations and Rely upon Different Aneuploid Chromosomes. MBio. 2023;14: e00227–23.

76. Pompei S, Lagomarsino MC. A fitness trade-off explains the early fate of yeast aneuploids with chromosome gains. Proceedings of the National Academy of Sciences. 2023;120: e2211687120.

77. Taylor AM, Shih J, Ha G, Gao GF, Zhang X, Berger AC, et al. Genomic and Functional Approaches to Understanding Cancer Aneuploidy. Cancer Cell. 2018;33: 676–689.e3.

78. Kolodner RD, Cleveland DW, Putnam CD. Cancer. Aneuploidy drives a mutator phenotype in cancer. Science. 2011. pp. 942–943.

79. Yona AH, Manor YS, Herbst RH, Romano GH, Mitchell A, Kupiec M, et al. Chromosomal duplication is a transient evolutionary solution to stress. Proc Natl Acad Sci U S A. 2012;109: 21010–21015.

80. Hirakawa MP, Martinez DA, Sakthikumar S, Anderson MZ, Berlin A, Gujja S, et al. Genetic and phenotypic intra-species variation in Candida albicans. Genome Res. 2015;25: 413–425.

81. Flowers SA, Barker KS, Berkow EL, Toner G, Chadwick SG, Gygax SE, et al. Gain-of-function mutations in UPC2 are a frequent cause of ERG11 upregulation in azole-resistant clinical isolates of Candida albicans. Eukaryot Cell. 2012;11: 1289–1299.

82. Flowers SA, Colón B, Whaley SG, Schuler MA, Rogers PD. Contribution of clinically derived mutations in ERG11 to azole resistance in Candida albicans. Antimicrob Agents Chemother. 2015;59: 450–460.

83. Rybak Jeffrey M., Sharma Cheshta, Doorley Laura A., Barker Katherine S., Palmer Glen E., Rogers P. David. Delineation of the Direct Contribution of Candida auris ERG11 Mutations to Clinical Triazole Resistance. Microbiology Spectrum. 2021;9: e01585–21.

84. Burrack LS, Todd RT, Soisangwan N, Wiederhold NP, Selmecki A. Genomic Diversity across Candida auris Clinical Isolates Shapes Rapid Development of Antifungal Resistance In Vitro and In Vivo. MBio. 2022;13: e0084222.

85. Li J, Aubry L, Brandalise D, Coste AT, Sanglard D, Lamoth F. Upc2-mediated mechanisms of azole resistance in Candida auris. Microbiol Spectr. 2024; e0352623.

86. Glazier Virginia E., Kramara Juraj, Ollinger Tomye, Solis Norma V., Zarnowski Robert, Wakade Rohan S., et al. The Candida albicans reference strain SC5314 contains a rare, dominant allele of the transcription factor Rob1 that modulates filamentation, biofilm formation, and oral commensalism. MBio. 2023;14: e01521–23.

87. Vande Zande P, Siddiq MA, Hodgins-Davis A, Kim L, Wittkopp PJ. Active compensation for changes in TDH3 expression mediated by direct regulators of TDH3 in Saccharomyces cerevisiae. PLoS Genet. 2023;19: e1011078.

88. Shen J, Guo W, Köhler JR. CaNAT1, a heterologous dominant selectable marker for transformation of Candida albicans and other pathogenic Candida species. Infect Immun. 2005;73: 1239–1242.

89. Veri AO, Miao Z, Shapiro RS, Tebbji F, O’Meara TR, Kim SH, et al. Tuning Hsf1 levels drives distinct fungal morphogenetic programs with depletion impairing Hsp90 function and overexpression expanding the target space. PLoS Genet. 2018;14: e1007270.

90. Gerami-Nejad M, Forche A, McClellan M, Berman J. Analysis of protein function in clinical C. albicans isolates. Yeast. 2012;29: 303–309.

91. Patro R, Duggal G, Love MI, Irizarry RA, Kingsford C. Salmon provides fast and bias-aware quantification of transcript expression. Nat Methods. 2017;14: 417–419.

92. Soneson C, Love MI, Robinson MD. Differential analyses for RNA-seq: transcript-level estimates improve gene-level inferences. F1000Res. 2015;4: 1521.

93. Love MI, Huber W, Anders S. Moderated estimation of fold change and dispersion for RNA-seq data with DESeq2. Genome Biol. 2014;15: 550.

94. Skrzypek MS, Binkley J, Binkley G, Miyasato SR, Simison M, Sherlock G. The Candida Genome Database (CGD): incorporation of Assembly 22, systematic identifiers and visualization of high throughput sequencing data. Nucleic Acids Res. 2017;45: D592–D596.

95. Todd RT, Braverman AL, Selmecki A. Flow Cytometry Analysis of Fungal Ploidy. Curr Protoc Microbiol. 2018;50: e58.

96. Selmecki A, Forche A, Berman J. Aneuploidy and isochromosome formation in drug-resistant Candida albicans. Science. 2006;313: 367–370.

97. Bushnell B. BBTools software package. 2014. Available online: http://sourceforgenet/projects/bbmap.

98. Li H. Aligning sequence reads, clone sequences and assembly contigs with BWA-MEM. arXiv [q-bio.GN]. 2013. Available: http://arxiv.org/abs/1303.3997

99. Danecek P, Bonfield JK, Liddle J, Marshall J, Ohan V, Pollard MO, et al. Twelve years of SAMtools and BCFtools. Gigascience. 2021;10. doi:10.1093/gigascience/giab008

100. Andrews S. FastQC: A quality control analysis tool for high throughput sequencing data. Github; Available: https://github.com/s-andrews/FastQC

101. Okonechnikov K, Conesa A, García-Alcalde F. Qualimap 2: advanced multi-sample quality control for high-throughput sequencing data. Bioinformatics. 2016;32: 292–294.

102. Ewels P, Magnusson M, Lundin S, Käller M. MultiQC: summarize analysis results for multiple tools and samples in a single report. Bioinformatics. 2016;32: 3047–3048.

103. Abbey DA, Funt J, Lurie-Weinberger MN, Thompson DA, Regev A, Myers CL, et al. YMAP: a pipeline for visualization of copy number variation and loss of heterozygosity in eukaryotic pathogens. Genome Med. 2014;6: 100.

104. Van der Auwera GA, O’Connor BD. Genomics in the Cloud: Using Docker, GATK, and WDL in Terra. “O’Reilly Media, Inc.”; 2020.

105. Cingolani P, Platts A, Wang LL, Coon M, Nguyen T, Wang L, et al. A program for annotating and predicting the effects of single nucleotide polymorphisms, SnpEff: SNPs in the genome of Drosophila melanogaster strain w1118; iso-2; iso-3. Fly . 2012;6: 80–92.

106. Robinson JT, Thorvaldsdóttir H, Wenger AM, Zehir A, Mesirov JP. Variant Review with the Integrative Genomics Viewer. Cancer Res. 2017;77: e31–e34.

107. Hornby JM, Nickerson KW. Enhanced production of farnesol by Candida albicans treated with four azoles. Antimicrob Agents Chemother. 2004;48: 2305–2307.

